# Mapping breast cancer lineage in radiation and immunotherapy using the REMAP mouse

**DOI:** 10.64898/2026.04.15.718689

**Authors:** Abigail C. Marshall, Jackie Vahey, Meisam Bagheri, Rachel Saxe, Jennifer Fields, Fred Kolling, Aaron McKenna

**Affiliations:** Department of Molecular and Systems Biology, Geisel School of Medicine, Dartmouth College, Hanover, NH; Dartmouth Cancer Center, Lebanon, NH

## Abstract

We present REMAP (Recording Evolution in Mammary tumors via Active PyMT), a lineage-tracing mouse model that integrates inducible CRISPR recording with the MMTV-PyMT model of hormone receptor-positive (HR+) breast cancer. Inducible Cas9 drives editing of MARC1 homing guide RNAs (hgRNAs), generating heritable lineage marks, and enables reconstruction of clonal relationships. Using REMAP, we profiled tumor evolution and response to radiation combined with anti-PD1 immunotherapy. Treatment reduced tumor burden locally and systemically, and single-cell RNA sequencing revealed remodeling of the tumor microenvironment (TME). We identified metastatic clones present across primary tumors and distant sites, which exhibited elevated epithelial-mesenchymal transition (EMT) programs as a heritable clonal state. Treatment reduced EMT-associated transcriptional programs and reshaped immune composition, with radiation driving clonal expansion of T cells and reduced repertoire diversity. In contrast, cancer-associated fibroblast clones spanned multiple transcriptional states, indicating substantial stromal plasticity. Together, REMAP enables high-resolution coupling of clonal history and cellular state in vivo, revealing that tumor progression, metastasis, and therapeutic response are governed by heritable lineage programs.

## Introduction

Breast cancer is among the most common cancer types and is a leading cause of cancer mortality in women^1,2^. While improvements in screening and therapeutics have greatly improved outcomes, certain types of breast cancer remain challenging to treat. While primary HR+ breast cancer has a favorable prognosis with five-year survival exceeding 90%, patients remain at risk for recurrence even decades after initial diagnosis^3–5^, and treatment of recurrent disease can be difficult^6^.

Immunotherapy has shown promise in treating aggressive, primary HR- breast cancers, but has so far been less effective in more common ER-positive subtypes^7–9^. This has been attributed to HR+ tumors having a more “immune cold” TME^10–14^. Estrogen is believed to have an immunosuppressive effect by decreasing tissue-resident T cells, supporting suppressive regulatory immune cell types, and impairing immune cell antitumoral effects^7,14–18^. Recent trials of immunotherapy, specifically those targeting the PD-1/PD-L1 interaction, show benefits in HR+ breast cancer patients, suggesting that at least some of these “immune-cold” tumors can benefit from immune checkpoint blockade^9,14,19–21^. This has led to interest in approaches that remodel the TME immune-scape to promote immunotherapy’s efficacy in HR+ breast cancers^7,14,18,22,23^. One such approach utilizes neoadjuvant radiation to shift an “immune cold” TME towards a more immunostimulatory TME to counter estrogen’s immunosuppressive impact and augment immunotherapy^23,24^.

Accurate preclinical models are critical for assessing the effectiveness of any therapy and understanding a treatment’s impact on the TME and immune cell populations. Mouse models of breast cancer that rely on cell injection cannot faithfully replicate host-tumor factors that contribute to tumor development and an immunosuppressed TME. Instead, spontaneous tumor models are crucial to recapitulate the host-tumor interplay^14,25^. The most widely used spontaneous breast cancer murine model is the transgenic mouse mammary tumor virus (MMTV) polyomavirus middle T (PyMT) antigen model^26,27^. This model follows stages of progression that parallel those observed in breast cancer patients and leads to tumors that are transcriptionally similar to human HR+ breast cancer. During early stages tumors are strongly HR+ ^26–28^, but display heterogeneous HR levels at advanced stages, paralleling what is seen in some recurrent HR+ breast cancers^27–31^.

Initial reports of single-agent immunotherapy in the MMTV-PyMT model show slight tumor delay but no tumor regression, consistent with immunotherapy outcomes in HR+ patients. Combining anti-CTLA4 with radiation stagnated tumor growth but did not significantly alter immune cell type distribution in the TME^32^. These preliminary reports warrant more detailed investigation.

Cellular lineage tracing offers a powerful method to assess the impact of neoadjuvant radiation and immunotherapy on all cells in the TME. Lineage tracing enables the recording of cell relationships across cell divisions to shed light on clonal dynamics^33–36^. Recent work using Confetti, Cre-recombinase, or DRAG lineage systems to assess clonality in the MMTV-PyMT model demonstrated tumor clones can have varying paths along the EMT trajectory^37,38^. These works provide high-resolution insight into clonal dynamics driving cancer progression, but have yet to describe the clonal dynamics in the context of treatment. Additionally, monitoring the lineage of both non-malignant immune cells and malignant cells will be vital to understanding how TME cell populations are altered.

To achieve lineage tracing in all cell types in the MMTV-PyMT model, we employ the MARC1 system (**Fig. 1A**)^39,40^. The MARC1 mouse uses multiple hgRNAs integrated throughout the genome as dynamic barcodes that record cell relationships across all cell types. We use an inducible Cas9 (iCas9) mouse to begin lineage tracing during embryonic development to coincide with mammary tissue formation ^41^. By combining iCas9, MARC1, and MMTV-PyMT mice into a single line, we created an inducible lineage-tracing mouse model of spontaneous, immune-intact, HR+ breast cancer. We then used this model to compare typical tumor progression versus neoadjuvant radiation and anti-PD1 immunotherapy to uncover the effects of this paired treatment regimen in HR+ breast cancer.

**Fig. 1:**
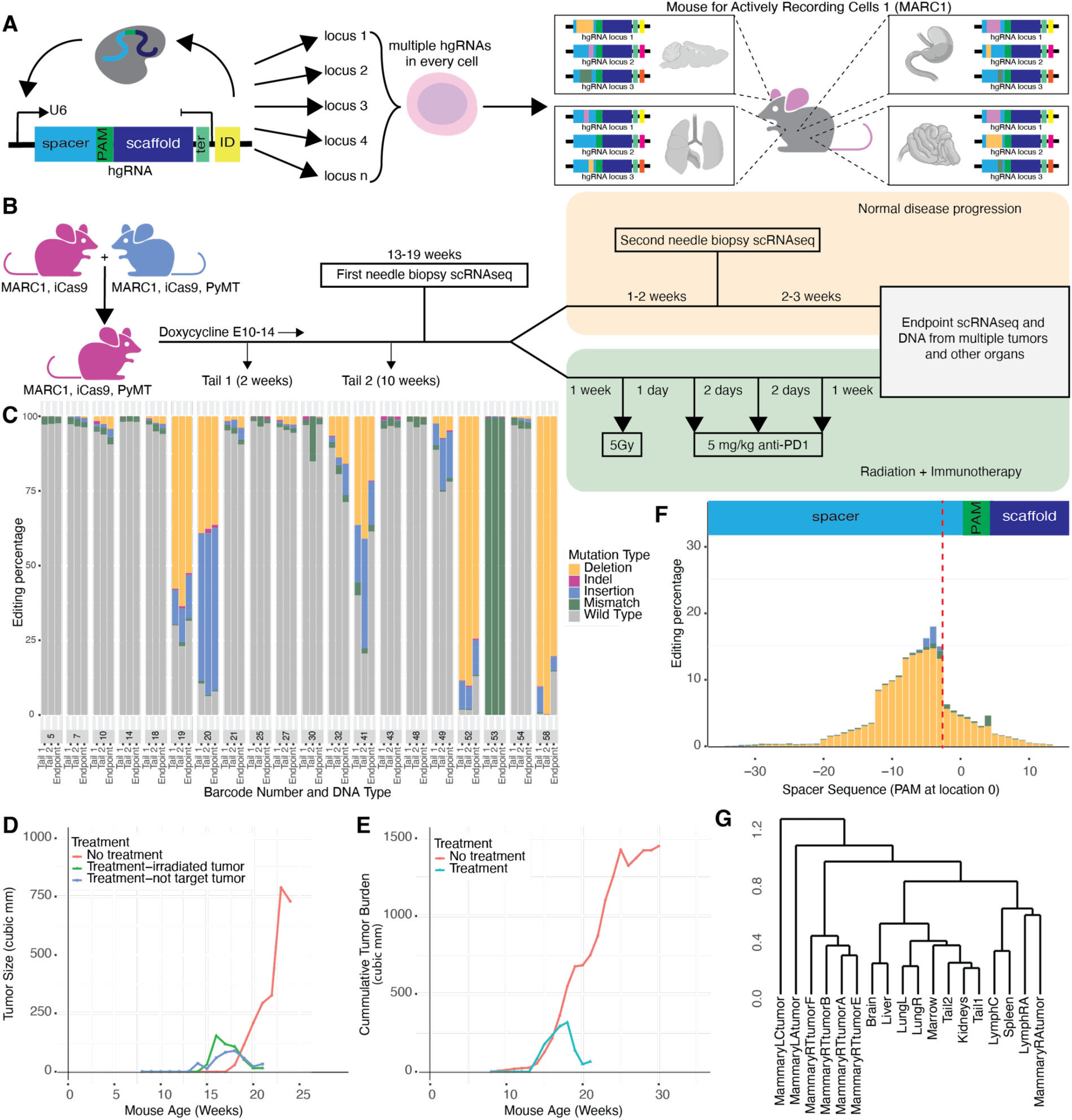
REMAP breast cancer lineage tracing. **A)** Schematic of REMAP mouse. **B)** Schematic of female REMAP mice and timeline of doxycycline exposure, tumor biopsies, radiation, anti-PD1 therapy, and endpoint sample collections for scRNAseq and DNA analyses. **C)** Percentage of hgRNA barcode reads from DNA with editing detected in 3 untreated and 3 treated REMAP mice grouped by sample type (tail1, tail 2, or endpoint sample). **D)** Percentage of edit types called in DNA sequencing at each position in hgRNA reads (expected Cas9 cut site indicated with red dashed line). **E)** Size of tumors in REMAP mice that received no treatment (red), tumors in treated mice that were not targeted with radiation (blue), and tumors in treated mice that were irradiated (green). **F)** Cumulative tumor burden in untreated (red) and treated (blue) REMAP mice. **G)** Example dendrogram (from mouse F221) constructed using DNA editing patterns to group tissue types by similarity of editing outcomes across all hgRNA integrations.

## Results

To create a lineage tracing model of HR+ breast cancer, we interbred Mouse for Actively Recording Cells 1 (MARC1) founder mice with an iCas9 line^39,42^. This was then crossed with the MMTV-PyMT mouse line to create our REMAP mouse line (**R**ecording **E**volution in **M**ammary tumors via **A**ctive **P**yMT) (**Fig. 1B**). Female REMAP mice were exposed to doxycycline beginning in utero to trigger Cas9 expression and enable editing of MARC1 hgRNAs. Editing of hgRNAs was assessed at two weeks and ten weeks of age by DNA extracted from tail clips and with DNA sampled from organs and tissues at endpoint (**Fig. 1C, Supplemental Table 1**).

REMAP mice were then separated into treatment and control arms (n=3). Mice developed multiple mammary gland tumors on a timeline consistent with previous reports ^262,263^. We performed serial mammary gland needle biopsies to evaluate the tumor composition and profile the TME^43^ (**Fig. 1B**). Three mice then received 5Gy radiation one week after biopsy and three doses of IP 5mg/kg anti-PD1 following radiation (**Fig. 1B**). Endpoint in these mice was defined as one week after the final anti-PD1 dose. The cumulative tumor burden decreased with treatment, with regression not only in tumors that were irradiated but also in other, nonirradiated tumors within the treated mice (**Fig. 1D,E**).

We then sequenced lineage recorders from both the endpoint tumor and other organs from all 6 mice. We detected active lineage recording in the expected region of the hgRNAs, around 3-5 nucleotides upstream of the PAM (**Fig. 1F**). Our inducible system also created more diverse lineage recording outcomes when compared to a constitutive-Cas9 mouse (**Supplemental Fig. 1A,B**), and REMAP mice without Cas9 or doxycycline had low background levels of CRISPR editing, outside of germline changes in two hgRNAs (**Supplemental Fig. 1C,D**).

Across all samples, we identified 191,541 unique lineage outcomes (**Supplemental Fig. 3A**). Of these, 47,641 were found in more than one mouse; this recurrent editing occurred at a much higher rate than in previously published work, likely due to our longer experimental timeline which may allow hgRNAs to edit to more common, terminal states^39,40^. Deletions were the most prevalent editing outcome, consistent with previous reports (**Supplemental Fig. 2, Supplemental Table 2)**^39,40^. Within each REMAP mouse, we used patterns of shared edits across tissue types to infer the germline-relatedness of bulk tissues (**Fig. 1G, Supplemental Fig. 4**).

We then collected single-cell RNA-seq profiles for 189,468 cells across the biopsies and endpoint tumors from three untreated and three treated mice (**Fig. 2A**). We used marker gene expression to annotate cells, resulting in cell type labels that cluster tightly on the integrated gene expression UMAP, across mice, timepoints, and sample (**Supplemental Fig. 5A-C**) ^44^.

**Fig. 2:**
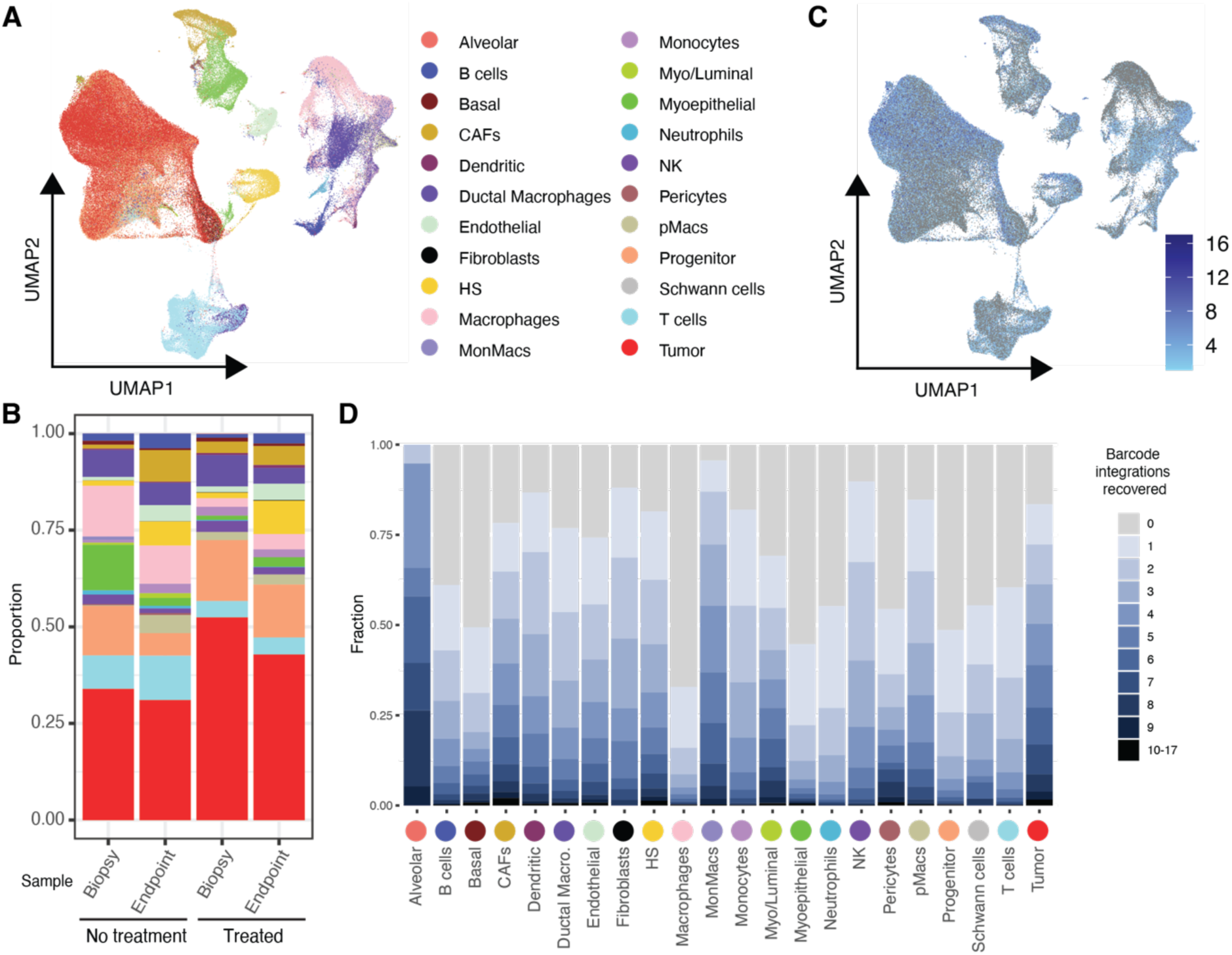
Single-cell profiling and recovery of REMAP recorders. **A)** UMAP of all cells collected across all timepoints in 3 untreated and 3 treated REMAP mice, colored by cell type annotation. **B)** Cell type proportions from biopsy (first biopsy for untreated mice) and biopsied tumor endpoint material grouped by untreated or treated mice, colors correspond to key in panel A. **C)** UMAP colored by the number of hgRNA barcodes recovered per cell. **D)** Proportion of hgRNAs recovered per cell type.

We detected many expected epithelial cell types: malignant tumor, alveolar, basal, hormone sensing (HS), myoepithelial, myoluminal, and progenitor cells. We also identified stromal cell types, including fibroblasts, cancer-associated fibroblasts (CAFs), endothelial, pericytes, and Schwann cells. Diverse immune cell populations were present, including B cells, T cells, dendritic cells, neutrophils, natural killer (NK) cells, monocytes, and various subtypes of macrophages. In untreated mice, no consistent pattern of cell proportion shifts emerged across mice with tumor progression (**Fig. 2B**, **Supplemental Fig. 5D**). In all treated mice, tumor cell proportion decreased from biopsy to endpoint, consistent with tumor volume regression. The fraction of HS, myoepithelial, and CAFs increased across all treated mice (**Fig. 2B**, **Supplemental Fig. 5D**).

We adapted an approach similar to that used by Islam et al to capture transcribed MARC1 lineage recorders (see Methods, **Supplemental Fig. 6**)^45^. Due to an upstream terminator sequence, hgRNA transcripts recovered from RNA do not contain the static identifiers marking each unique hgRNA integration^39,40,45^. This complicates analysis of edited hgRNA as heavily edited hgRNA sequences from different barcode integrations may converge. To overcome this challenge and accurately attribute edits from single cell sequencing, we utilized the edited hgRNAs from DNA sequencing to guide our interpretation of scRNAseq data (see Methods**, Supplemental Fig. 7**). Using this approach, we confidently identified 144,428 (43.2%) of the 334,417 edits detected in DNA. The editing patterns from scRNAseq aligned well with patterns from DNA sequencing (**Supplemental Fig. 8A,B**) and varied across cell types (**Fig. 2C, D, Supplemental Fig. 8C**). Recovery varied by cell type, and our capture of recorders in tumor cells was higher than in somatic cells (**Fig. 2D**). Cell type relationships inferred by sparse patterns of shared scRNAseq hgRNAs highlighted shared ancestry among tumor cells and among immune cell populations (**Supplemental Fig. 9**).

To explore progenitor and tumor cell populations in more detail, we integrated data from untreated and treated mice, selected progenitor and tumor cells, and re-clustered this subset of cells (**Fig. 3A-C, Supplemental Table 4**). The progenitor and tumor cells exhibited transcriptional heterogeneity, separating into ten clusters (**Fig. 3A**). This diversity is consistent with previous reports using the MMTV-PyMT model and intratumoral heterogeneity observed in breast cancer patient samples. Clusters 3 and 4 were predominantly tumor cells, Clusters 5 and 6 contained mostly progenitor cells, and other clusters contained a mix of cell types (**Fig. 3B**).

**Fig. 3:**
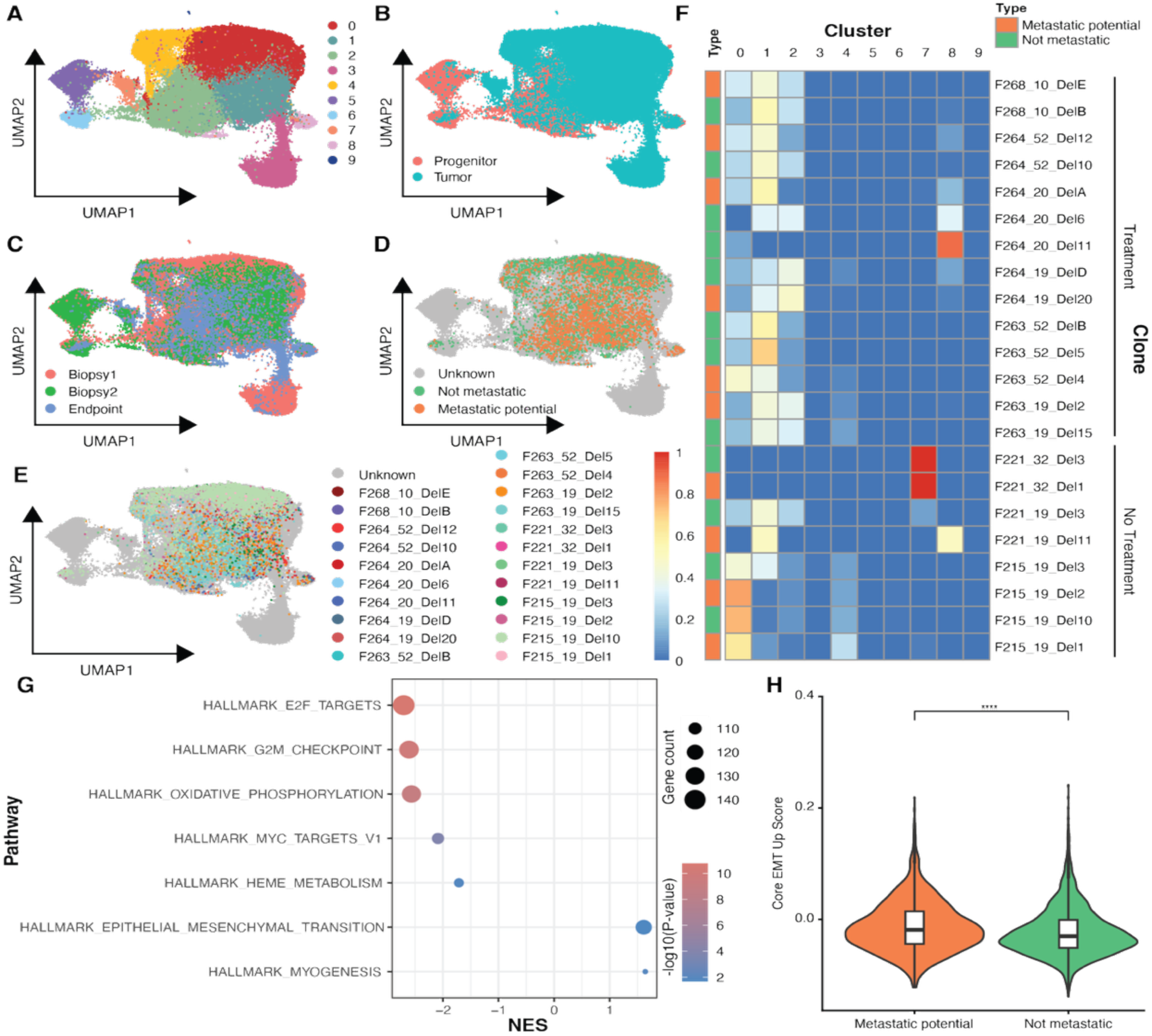
Effect of radiation and immunotherapy on REMAP mice. A-C) UMAPs of tumor and progenitor cells from 3 untreated and 3 treated REMAP mice colored by cluster assignment (**A**), cell annotation (**B**), or timepoint of collection (**C**). **D)** UMAP colored by REMAP clone type of metastatic potential (orange) or not metastatic (green). **E)** UMAP colored by REMAP clone identity; red, orange, and pink clones have metastatic potential while green and blue clones are not metastatic clones. **F)** Cluster distributions of REMAP clones from panel E showing proportion of each clone assigned to each cluster. **G)** Gene set enrichment analysis of hallmark pathways showing significantly upregulated or downregulated pathways in cells of clones with metastatic potential versus not metastatic. **H)** Core EMT Up Score in cells of clones with metastatic potential or not.

Across all clusters, markers associated with luminal B and HER2-positive breast cancer were higher than luminal A or TNBC markers (**Supplemental Fig. 10A**), consistent with the phenotypic shift from luminal B-like to HER2-positive-like observed in later stages of the MMTV-PyMT model^28,46^. The breast cancer stem cell gene signature reported by Jiang et al was highest in Clusters 1, 7 and 8^47^. Applying transcriptional states defined by Yeo et al, Cluster 3 appeared to be a mix of luminal progenitor involuting (LP-INV) and luminal progenitors (LP), whereas Clusters 4 and 5 aligned with the alveolar-primed state^48^. Cluster 3 exhibited higher expression of mesenchymal genes *Sparc*, *Vim*, and *Col1a2*, suggesting tumor cells within this cluster are less epithelial and shifted towards a hybrid EMT state. Cluster 4 was defined by proliferative markers such as *Birc5* and *Mki67* (**Supplemental Fig. 10A, Supplemental Table 4**).

We observed a number of hgRNA mutations shared between the tumor single-cell populations and tissue DNA samples from the lungs and/or lymph nodes of the same mouse (**Supplemental Fig. 11)**. We hypothesized that these hgRNA edits marked sites where metastasis had occurred, common in the MMTV-PyMT model. Using these shared hgRNA edits, we identified 10 metastatic clones encompassing 2,245 cells (**Fig. 3D,E, Supplemental Table 3**). We also created a set of clones where we could exclude metastasis in the mouse, labeling 12 non-metastatic clones with 4,677 cells. These clones clustered spatially on the UMAP and were predominantly present in one to three clusters (**Fig. 3E,F**).

Differential gene expression revealed transcriptomic differences between metastatic and non-metastatic clones (**Supplemental Fig. 12A, Supplemental Table 5**). Gene set enrichment analysis (GSEA) using hallmark pathways identified EMT as upregulated in potentially metastatic clones. Conversely, growth-related pathways, oxidative phosphorylation, and *Myc* targets were downregulated in metastatic clones (**Fig. 3G**). This suggests that clones with metastatic potential have increased expression of EMT factors on a clonal level, priming these clones for metastasis. We further validated this by creating a “Core EMT Up Score” derived from expression of the upregulated EMT genes in the core breast cancer EMT signature described by Taube et al^49^. Cells of the metastatic clones had significantly higher Core EMT Up Scores than cells of non-metastatic clones (adjusted p-value = 1.2e-20 **Fig. 3H**).

To assess the impact of treatment on all tumor and progenitor cells, we pseudobulked tumor cells by sample and conducted differential gene expression and GSEA. There were significant transcriptional changes between pre-treatment biopsies and post-treatment endpoint tumor cells (**Supplemental Fig. 12B**) and between treated and untreated endpoint tumor cells (**Supplemental Fig. 12C**). We also found transcriptional changes in treated endpoint cells from targeted, irradiated tumors compared to those that did not receive direct irradiation (**Supplemental Fig. 12D**). Collectively these comparisons revealed the impact of radiation, anti-PD, and their combination on cancer cells, notably including a decrease in EMT pathway expression following combination therapy.

As our lineage recordings spanned tumor and somatic tissues, we next focused on immune components of the tumor microenvironment, including CAFs and T cells. To analyze clonal state distributions of CAFs, we re-clustered only fibroblasts and CAFs and annotated their transcriptional states using the subtypes described by Bartoschek et al (**Fig. 4A**)^50^. Consistent with Bartoschek et al, we observed a slight increase in vascular CAF (vCAF) proportion with tumor progression regardless of treatment, however matrix CAF (mCAF) fraction was relatively stable across time and treatment (**Supplemental Fig. 13A,B**). We then identified 50 distinct CAF clones by hgRNA edits, encompassing 2,360 cells (**Fig. 4B, Supplemental Fig. 13C, Supplemental Table 6**). Interestingly, all clones contained cells of multiple subtypes, suggesting less stringent origins and plasticity in CAF states (**Fig. 4C**).

**Fig. 4:**
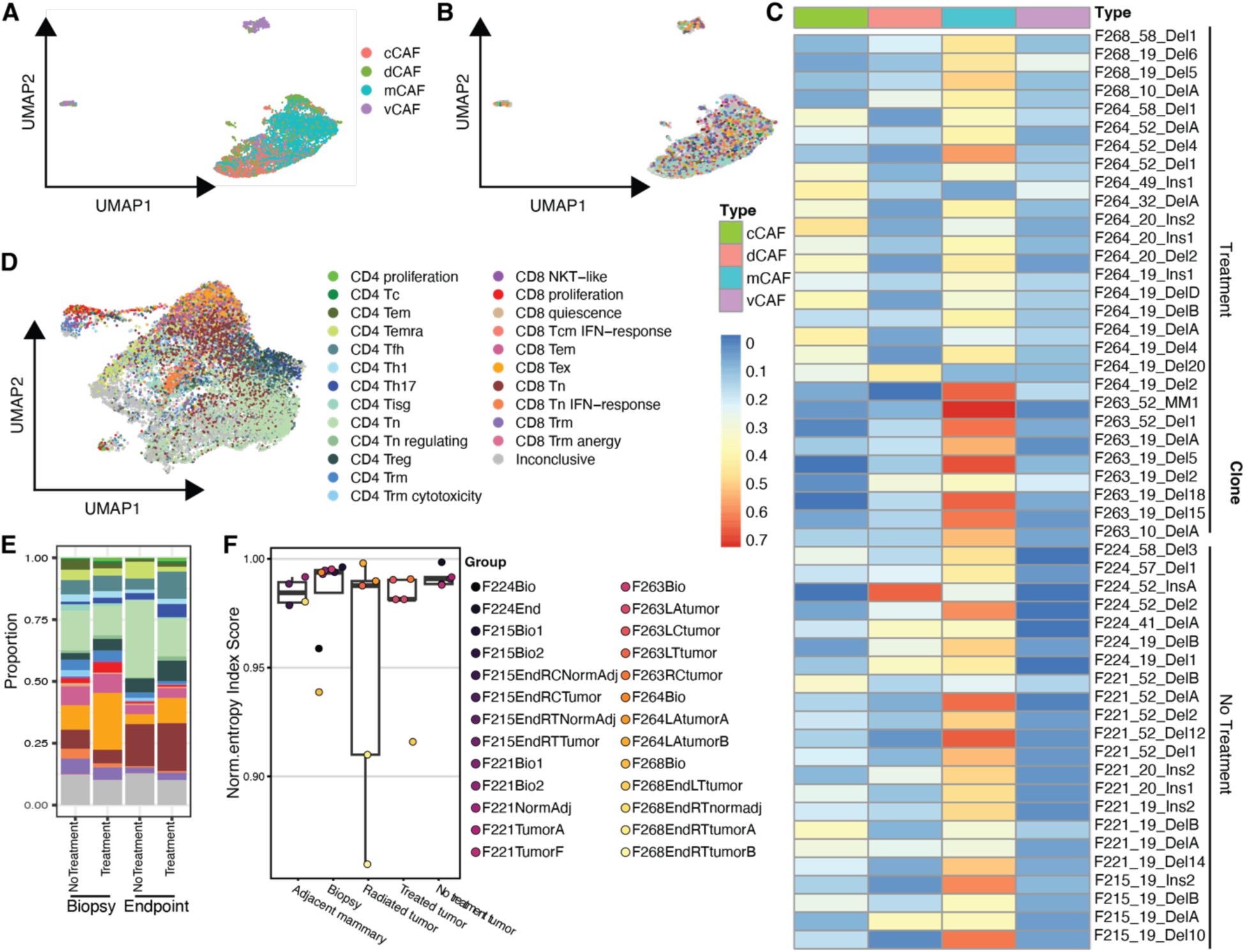
Immune cell clones modulated by treatment. A,B) UMAP of CAFs and fibroblasts from 3 untreated and 3 treated REMAP mice colored by CAF subtype (**A**) or REMAP clone designation (**B**). **C)** Cluster distributions of REMAP clones from panel B showing proportion of each clone assigned to each CAF subtype. **D)** UMAP of T cells from 3 untreated and 3 treated REMAP mice colored by T cell subtype annotation. **E)** Proportion of T cell subtypes in treated and untreated biopsy (first biopsy for untreated mice) and endpoint samples, colors correspond to key in panel D. **F)** Normalized entropy score for T cell clone library grouped by sample type, showing sample means and confidence interval from bootstrapping analysis. Sample types include adjacent mammary tissue collected at endpoint; biopsy material from pre-treatment and untreated tumors; radiated tumors in treated mice that received anti-PD1; non-irradiated tumors in treated mice that received anti-PD1; and untreated tumors at endpoint.

Next, we isolated and re-clustered T cells and utilized STCAT to annotate subtypes (**Fig. 4D**)^51^. We then assessed distributions of T cell subtypes in biopsy and endpoint samples from treated and untreated mice (**Fig. 4E, Supplemental Fig. 14A**). There was a shift in CD4+/CD8+ balance towards more CD4+ T cells with time regardless of treatment. In treated mice, follicular helper T cells notably expanded. This T cell subtype plays a key role in tertiary lymphoid structures and crosstalk with B cells, illustrating a potential role for B cells in immunotherapy response. Proliferative CD8+ T cells were detected at a lower level in endpoint samples relative to biopsies. Sparse recovery of hgRNAs in T cells restricted our ability to use edited barcodes to define T cell clones, however T cells record their clonal identities via V(D)J sequence recombination of the T cell receptors (TCR) (**Supplemental Fig. 14B**). We utilized TRUST4 to extract V(D)J sequences from our scRNAseq data and used scRepertoire to label T cell clones by the amino acid sequence of their TCRs^52,53^. Diversity analysis of T cell clones by sample type revealed a clear impact of radiation (**Fig. 4F**). There was lower normalized diversity of T cell clones in radiated tumors, with an expansion of T cell clones responding to damaged tumor cells. Anti-PD1 alone did not have a strong influence on T cell clone diversity, highlighting the importance of neoadjuvant radiation to boost immunotherapy response in this ER-positive breast cancer model.

## Discussion

Here we present the REMAP mouse and utilize it to chart the development and evolution of breast cancer under the selective pressure of treatment. REMAP allows researchers to track the spontaneous emergence of tumor cells from somatic tissues, enabling the longitudinal, multi-tissue lineage tracing in an immunocompetent cancer model. We show that an inducible Cas9 system can record lineage over developmental time, recording relationships between organs and tissues, and tracking the emergence of tumor cells. Using both DNA and RNA recovery, we show that editing is highly variable across integrated recorders throughout time and generates unique marks labeling tumor formation after weeks of development.

With REMAP, we observed enrichment of canonical EMT signatures, linked to clonal populations evolved to seed metastatic sites. These signatures were enriched within the primary tumor, suggesting that this EMT state is heritable and evolved clonally. This supports a model where metastatic potential is an intrinsic, lineage-linked state rather than solely induced by the metastatic niche. This contrasts with CAFs, where individual clones span multiple CAF states.

This suggests that CAF identity is plastic and not strictly lineage-determined, whereas tumor cells rapidly evolve to a stable clonal state.

Treatment using radiation and immunotherapy reduced tumor burden across all tumors, including tumors not directly targeted by radiation. This suggests the combination of radiation and anti-PD1 can lead to local tumor control, as well as control of other distal tumors within the mouse. The regression of nonirradiated tumors may be due to the effects of anti-PD1 alone or in conjunction with the abscopal effect of radiation. Our findings are consistent with previous work and lend support to the addition of neoadjuvant radiation to prime ER-positive tumors to respond better to immune checkpoint blockage^9,23,32,54^. We also observed reduced TCR diversity after radiation, with expansion of specific clones. Radiation acts here as a key initiator of adaptive immune responses, though we were unable to fully disentangle the roles of immunotherapy and radiation.

There are limitations to the REMAP approach. Lineage recording was variable and generally confined to a limited number of recorders integrated into the genome, consistent with previous reports^39^. Compared with a constitutive Cas9 system, our inducible system exhibited greater variability in editing outcomes, though we still observed low editing rates at many sites. Capture rates from single-cell sequencing certainly could also be improved, especially for non-tumor cells, possibly using targeted approaches or in vitro transcription approaches^55^. Our analysis pipeline took a more conservative approach than previous efforts to overcome the lack of transcribed integration markers, leveraging matched events observed in DNA sequencing across organs and tissues. Lastly, we are drawing inferences from single-cell lineage tracing with a very limited number of mice, and our approach has limited power to disentangle the effects of the immunotherapy and radiation on our tumor regression or immune cell dynamics.

Together, these results establish REMAP as a powerful platform for linking lineage history with cellular state and therapeutic response in vivo, revealing how heritable tumor programs and environmentally driven microenvironmental dynamics jointly shape cancer evolution. By demonstrating that metastatic potential is encoded within clonal tumor lineages, while stromal compartments remain plastic, and that radiation can reprogram systemic immune responses to enhance tumor control, our study provides a framework for understanding how intrinsic and extrinsic forces interact during disease progression. Although further refinement of recording capacity, capture efficiency, and cohort size will be important, REMAP opens the door to mechanistic studies of tumor development in autochthonous mice. REMAP, and other approaches, will allow us to dissect clonal evolution, offering a path toward identifying lineage-informed vulnerabilities that can be exploited to improve therapeutic strategies^38^.

## Methods

### Ethics approval

All animal experiments were conducted in accordance with the guidelines of and with the approval of the Institutional Animal Care and Use Committee at Dartmouth College.

### Animals

Mice were maintained in pathogen-free housing at the Center for Comparative Medicine and Research in Dartmouth Hitchcock Medical Center. MARC1-PB7 (RRID: MMRRC_065424-UCD) mice were obtained from University of California Davis Mutant Mouse Resource and Research center. KH2/iCas9 mice (Strain #:029415, RRID:IMSR_JAX:029415) were a gift from the Huang lab at Dartmouth. Constitutive Cas9 (Strain #:026179 RRID:IMSR_JAX:026179) and PyMT lines (Strain#:022974, RRID:IMSR_JAX:022974) were obtained from The Jackson Laboratory.

MARC1 and iCas9 mice were crossed to create the MARC1-iCas9 line, which was then bred with PyMT males to generate MARC1-iCas9-PyMT males. MARC1-iCas9-PyMT males were bred with MARC1-iCas9 females to produce experimental, iCas9 mice. For constitutive Cas9 experimental mice, Cas9 females were bred with MARC1-iCas9-PyMT males. All mice were genotyped for relevant loci following The Jackson Laboratory’s protocols, mice bred to produce experimental mice were also genotyped for MARC1 hgRNAs as previously described^39,40^.

For iCas9 activation (via doxycycline) to edit MARC1 hgRNAs, MARC1-iCas9 females bred to MARC1-iCas9-PyMT males were closely monitored with daily weight checks. When a female gained at least 2g within a 7-day period she was suspected to be pregnant, with E0 considered to be 7 days prior, and removed from the breeding cage^56^. Between E10-E14 to trigger hgRNA editing in utero, pregnant dames were switched to 2mg/kg doxycycline in drinking water to induce iCas9 expression. Doxycycline exposure in water was continued. Embryonic days were confirmed based on when pups were born. At two weeks of age, female pups from experimental litters were tailed for genotyping and weekly intraperitoneal (IP) injections of 15mg/kg doxycycline were started to further improve iCas9 expression. Female pups with correct genotypes were raised as experimental mice with continuous 2mg/ml doxycycline in drinking water and weekly IP 15mg/kg doxycycline injections. Experimental mice were re-tailed at 10 weeks of age to assess hgRNA editing patterns. Experimental mice were closely monitored for mammary tumor growth. Tumor size was measured at least three times a week with electronic calipers. Endpoint cumulative tumor burden was 1500mm^3^.

### Mammary gland biopsy

Once a tumor was palpable (>50mm^3^, mice aged 15-19 weeks), mice were anesthetized with isoflurane. A small incision was made, and the tumor was moved toward the incision to be more accessible. A 20G needle on a 3ml syringe was loaded with ∼500ul cold RPMI and negative pressure was established by slightly drawing back on the plunger. The needle was inserted into the tumor and rotated, all while negative pressure was maintained. Some of the RPMI was then flushed from the needle into a 1.7ml tube on ice to collect cells. The process of inserting, rotation, and flushing the needle was repeated at different points in the tumor to ensure collection of diverse tumor regions. After, the needle and syringe were flushed with more RPMI to ensure all cells were collected. Collected cells were kept on ice until single cell preparation started below. Incisions were closed with sutures and mice were closely monitored for wound healing. For repeat biopsies, the procedure outlined above was repeated at least 7 days after the first biopsy.

### Treatment regimen

One week after mammary gland biopsy, mice received 5Gy of radiation delivered via X-Rad 320 (Precision X-Ray Irradiation). Briefly, mice were anesthetized with isoflurane and positioned to expose the mammary tumor previously biopsied. Once mice were positioned and secured, a dynamic collimator was used to center the beam directly on the tumor and minimize the size of the field. Radiation was delivered with the following settings: 320kV, 12.5mA, SSD 40cm. The day after radiation, mice received the first of three doses of IP 5mg/kg anti-PD1 (bioXcell Clone RMP1-14, Cat# BE0146). Doses were given every other day. Endpoint was one week after the final anti-PD1 injection.

### Endpoint tumor and organ collection

When mice reached endpoint, determined by tumor burden for non-treated mice or time since treatment for treated mice, they were euthanized with CO2. All tissues collected at endpoint were either flash frozen on dry ice for subsequent gDNA extraction or placed into gentleMACS C Tubes (Miltenyi Cat# 130-093-237) on ice for subsequent single cell dissociation. Lymph nodes were dissected out first as they were easier to isolate before other tissues were disturbed. Then, mammary gland tumors were removed. Tumors were divided into different sections using razor blades, with some sections flash frozen and some used for single cell dissociation and sequencing. Tumor sections processed for single cell sequencing were further cut into smaller pieces before being placed in gentleMACS C Tubes (Miltenyi Cat# 130-093-237). Normal appearing mammary gland tissue adjacent to tumors was similarly collected. Next, any mammary glands without evident tumors were collected and flash frozen. After all mammary tissue was removed, other internal organs were dissected and flash frozen. To collect bone marrow, an 18G needle was used to make a hole in the bottom of a 0.5ml tube. Both femurs were placed in the 0.5ml tube, which was then placed in a 1.7ml tube, and centrifuged at > 10,000g for 30 seconds. The isolated bone marrow was then flash frozen.

### Single cell dissociation and sequencing

Material collected from mammary gland biopsies were moved to gentleMACS C Tubes (Miltenyi Cat# 130-093-237). Tumor and mammary gland material collected at endpoint were already collected in gentleMACS C Tubes. The 10X protocol “Tumor Dissociation for Single Cell RNA Sequencing” (10x CG000147, Rev B) was followed using the mouse tumor dissociation kit (Miltenyi Cat# 130-096-730). Once samples were mixed with dissociation enzymes in RPMI, they were mechanically digested using a gentleMACS Octo Dissociator with Heaters (Miltenyi Cat# 130-096-427) with the “37C_m_TDK_2” program. Cells were then filtered through a 40μM filter using RPMI to flush the strainer. For biopsy samples, due to much smaller numbers of cells, reduced volumes were used for resuspension, filtering, and subsequent steps. Next, red blood cell lysis was carried out using Miltenyi’s red blood cell lysis solution (Cat# 130-094-183). Samples were incubated with red blood cell lysis solution for five minutes at 4C.

Once tissues were dissociated, a sample of each resuspension was visually inspected with a microscope to quantify cells and live-dead cell proportions. If a sample contained >50% dead cells, dead cell removal was conducted according to 10x protocol “Removal of Dead Cells from Single Cell Suspensions for Single Cell RNA Sequencing” using Miltenyi dead cell removal kit (Cat# 130-090-101), LS columns (Cat# 130-042-401), and QuadroMACS separator (Cat# 130-090-976). Final cell resuspensions were prepared at ∼1000 cells/ul in 0.04% BSA.

A 10x Chromium X was utilized to capture single cells. For biopsy samples, 10x GEM-X Universal 5’ Gene Expression v3 reagents and Chromium GEM-X Single Cell 5’ Chip Kit v3 were used. For endpoint, four samples per mouse were multiplexed using 10x GEM-X Universal 5’ Gene Expression v3 4-plex reagents and GEM-X OCM 5’ Chip Kit v3 4-plex. Gene expression libraries were constructed following 10x protocol with two modifications. First, a custom capture primer (Supplemental Table 7) was spiked into the reverse transcription and cDNA amplification reactions. This reverse primer is complementary to the guide scaffold in the MARC1 hgRNA integrations and thus will enrich all MARC1 hgRNAs regardless of editing pattern. Second, half of the cDNA was taken before SPRIselect bead cleanup. This portion was used for MARC1 single cell hgRNA library preparation outlined below. This material was taken before SPRIselect cleanup to avoid loss of small, heavily edited hgRNAs. The remaining cDNA was processed for gene expression libraries according to 10X protocol. TCR libraries were prepared using 10x Mouse T Cell Mix 1 v2 (PN-2000256) following manufacturer’s protocol.

### Genomic DNA library preparation

All DNA was extracted using the DNeasy kit (Qiagen Cat# 69504) following the manufacturer’s protocol. Fresh tissue (tail clips) or small pieces of frozen tissue (endpoint samples) were digested overnight at 56C according to the Qiagen protocol. Once DNA was isolated, libraries were prepared using the primers from Leeper et al 2021^40^. For initial amplification, SBS3-PBLib-F and SBS9-PBLib-R primers (Supplemental Table 7) were used and the PCR reaction was stopped in the mid-exponential phase. Subsequent PCR reactions added Illumina p5 and p7 adaptors and Illumina P7 index to identify the sample.

### Single cell RNAseq MARC1 hgRNA library preparation

The portion of cDNA set aside before SPRIselect cleanup was cleaned with 2X ampure bead cleanup to retain DNA fragments of all sizes and avoid loss of heavily edited hgRNAs. The initial PCR amplification was conducted across eight replicates per sample using primers against TruSeq Read 1 and a portion of the scaffold sequence (Supplemental Table 7). Subsequent PCR reactions added TruSeq Read 2 indices and Illumina adaptors. Final libraries were assessed via TapeStation (Aligent Technologies) for size distribution and quantified via Qubit fluorometer and NEB NextQuant Library Quant kit (Cat # E7630L). Libraries were then diluted to the appropriate loading concentration for sequencing on an Illumina NextSeq2000.

### Sequencing and raw data processing

For bulk gDNA, TCR libraries, and single cell MARC1 hgRNA sequencing, bcl2fastq2 or BCL Convert (Illumina) was used to convert raw sequencing data to fastq format. For single cell RNA sequencing gene expression data, CellRanger mkfastq was used to convert raw sequencing data to fastq format. CellRanger count or CellRanger multi were used to call cells from non-multiplexed or on chip multiplexed experiments respectively.

### Cell type annotation

CellRanger-filtered matrices were analyzed using Seurat 5.3.0. Cells were filtered on number of genes and mitochondrial read percentage to remove low quality cells from each sample. The filtered matrices for biopsy and endpoint samples were merged for each mouse and normalized via Seurat’s NormalizeData(). Next, Seurat’s IntegrateLayers() with CCAIntegration was used to integrate data across different timepoints. Dimensional reduction and clustering (with resolution of two) was run on the integrated data to generate a UMAP for each mouse. The FindMarkers() function was utilized to generate a list of highest expressed genes for each cluster, this gene list was used to classify clusters as “immune-related” or “other.”

Next, the dataset for each mouse was subset into two separate datasets to split “immune-related” and “other” clusters. Dimensional reduction and clustering (with resolution of 0.1) was re-run on each subset to generate more precise clusters. These clusters were assigned cell types based on expression of marker genes as previously outlined^14,44^. The cell annotations from the subsets were then added as metadata to the full dataset to produce the cell type annotations for each mouse.

For CAF subtyping, Seurat’s AddModuleScore() was used to assign a score for each subtype using gene lists outlined by Bartoschek et al^50^. The subtype with the highest score was selected as the label for each cell.

For higher resolution T cell annotations, cells classified as T cells by the above approach were isolated from all mice and clustered together. Automated annotation was performed on this subset of data using STCAT^51^.

### TRUST4 and TCR analysis

Fastqs from gene expression and TCR sequencing were combined for each sample. To construct TCR sequences from the general and targeted sequencing, the combined data was processed with TRUST4^52^. Output from TRUST4 was then filtered to only cells that passed the gene expression quality filters outlined above. To focus on T cell dynamics, the filtered TRUST4 output was narrowed to only cells classified as αβ T cells by TRUST4. This dataset was then processed using scRepertoire^53^. T cell clonotypes were defined by the amino acid sequence of CDR3 regions determined by scRepertoire.

### MARC1 gDNA hgRNA identification and alignment

Fastqs from gDNA sequencing were processed using a custom analysis pipeline to identify edited MARC1 hgRNAs. BBMerge from the BBTools package was used to merge reads before sequences were extracted^57^. Relevant spacer sequences were identified using the barcode integration identifier and extracted sequences were aligned to their respective wild type sequence. The Bio.Align package was used for all alignments, with an open gap penalty of -25, a gap extension penalty of 0, a match score of 5, and a mismatch penalty of -4. These alignment penalties and scores were chosen to match those used in the AmpliCan tool for local alignments^58^. Each sample was processed separately to count wild type and edited MARC1 hgRNAs independently, then results were combined by mouse for analysis.

### MARC1 gDNA hgRNA whitelist generation

A whitelist of clearly identifiable edited hgRNAs was generated for each mouse through the DNA barcode identification pipeline. The whitelist requires that the edited hgRNA sequence be attributable to a single barcode integration identifier, that the edited hgRNA sequence be longer than 6 bases in length, and that the edited hgRNA sequence contains no Ns. If multiple identical edited hgRNA sequences were attributed to more than one barcode integration identifier and the difference between the frequency of the two top barcode integration identifiers was more than five percent, the sequence was allocated to the barcode integration identifier that appeared the most often. Edited hgRNA sequences that were attributed to more than one barcode integration identifier at similar frequencies were excluded from the whitelist. This whitelist was saved to be used in the single cell analysis pipeline and was generated independently for each mouse.

### MARC1 single cell RNA sequencing hgRNA identification and alignment

Fastqs from single cell RNA sequencing were processed using a custom analysis pipeline to identify edited MARC1 hgRNAs. Prospective edited hgRNA sequences, cell IDs, and molecule UMIs were extracted from the raw Fastq sequences. Prospective edited hgRNA sequences with cell IDs that were not found on the 10X cell ID whitelist were removed from analysis. Sequences with the same cell ID and UMI were merged to generate a single consensus sequence for that cell ID - UMI combination. After identifying consensus sequences containing wild type hgRNA sequences, the whitelist generated for each respective mouse was used to search through the remaining consensus sequences for edited hgRNA sequences. Edited hgRNA sequences were identified from each sample independently, then were combined by mouse for further analysis.

### Identification of potentially metastatic MARC1 clones and Core EMT Up Score

Edited hgRNA sequences that appear in at least 4% of reads from any one DNA sample or in at least 4% of cells from any one scRNAseq sample were named and further analyzed. To select potentially metastatic MARC1 hgRNA markers, we first bucketed DNA samples as possible metastatic sites (lymph node or lung samples), mammary (any mammary sample, including tumor and adjacent mammary), or other (kidney, liver, or tail samples). Then we identified edited hgRNAs that occurred in mammary and possible metastatic site samples, but not in other samples, using a 1% cutoff, as metastatic clonal barcodes. We also identified hgRNA identifier-matched non-metastatic edits, which were defined as occurring in only mammary samples, again using a 1% cutoff, with similar numbers of cells to their matching metastatic hgRNA. The matching barcodes were supplemented by edits that occurred across all sample types in cases where a matching non-metastatic labeled edit did not exist.

Seurat’s AddModuleScore was used to assign Core EMT Up scores based on expression of mouse orthologs of the genes identified as increasing along the breast cancer EMT spectrum in Taube et al’s core EMT signature^49^.

### Differential Gene Expression and Gene Set Enrichment Analysis

Differential gene expression was conducted comparing either groups of cells (i.e. potentially metastatic versus not metastatic clone cells) or pseudobulked values (for tumor cells bulked by sample, pseudobulked with Seurat’s AggregateExpression() function) using FindMarkers(). The resulting gene list was used for GSEA. GSEA was conducted with the fgsea package for hallmark pathways using default parameters^59^.

## Acknowledgements

We thank the members of the McKenna lab for discussion and support. Sequencing for the REMAP project was carried out in the Genomics and Molecular Biology Shared Resource (GMBSR, RRID:SCR_021293) at Dartmouth, which is supported by NCI Cancer Center Support Grant 5P30CA023108 and NIH S10 (1S10OD030242) awards. Single cell studies were conducted through the Dartmouth Center for Quantitative Biology in collaboration with the GMBSR with support from NIGMS (P20GM130454) and NIH S10 (S10OD025235) awards. Animal work was conducted in partnership with the Irradiation, Imaging, Microscopy & Animal Cancer Models (RRID:SCR_025077) at Dartmouth supported by Grant number P30CA023108. A.C.M. was supported by the Rosaline Borison Memorial Predoctoral Fellowship. This work was supported by DP2GM149750, and A.M. is supported by the Pew Biomedical Scholars program and the V Foundation.

## References

1. Arzanova, E. & Mayrovitz, H. N. The epidemiology of breast cancer. in Breast Cancer 1–20 (Exon Publications, 2022).

2. Amato, O., Guarneri, V. & Girardi, F. Epidemiology trends and progress in breast cancer survival: earlier diagnosis, new therapeutics. Curr. Opin. Oncol. 35, 612–619 (2023).

3. Calip, G. S. et al. Impact of time to distant recurrence on breast cancer-specific mortality in hormone receptor-positive breast cancer. Cancer Causes Control 33, 793–799 (2022).

4. Pedersen, R. N. et al. The incidence of breast cancer recurrence 10-32 years after primary diagnosis. J. Natl. Cancer Inst. 114, 391–399 (2022).

5. 5. Female Breast Cancer Subtypes - Cancer Stat Facts. *SEER* https://seer.cancer.gov/statfacts/html/breast-subtypes.html.

6. Lee, Y. J. et al. Prognosis according to the timing of recurrence in breast cancer. Ann. Surg. Treat. Res. 104, 1–9 (2023).

7. Dieci, M. V., Griguolo, G., Miglietta, F. & Guarneri, V. The immune system and hormone-receptor positive breast cancer: Is it really a dead end? Cancer Treat. Rev. 46, 9–19 (2016).

8. Goldberg, J. et al. Estrogen receptor mutations as novel targets for immunotherapy in metastatic estrogen receptor-positive breast cancer. Cancer Res. Commun. 4, 496–504 (2024).

9. Nader-Marta, G., Waks, A. G., Tolaney, S. M. & Mayer, E. L. Incorporating immunotherapy in the management of early-stage estrogen receptor-positive breast cancer. ESMO Open 9, 103977 (2024).

10. Dieci, M. V. et al. Update on tumor-infiltrating lymphocytes (TILs) in breast cancer, including recommendations to assess TILs in residual disease after neoadjuvant therapy and in carcinoma in situ: A report of the International Immuno-Oncology Biomarker Working Group on Breast Cancer. Semin. Cancer Biol. 52, 16–25 (2018).

11. Wagner, J. et al. A single-cell atlas of the tumor and immune ecosystem of human breast cancer. Cell 177, 1330–1345.e18 (2019).

12. Wu, S. Z. et al. A single-cell and spatially resolved atlas of human breast cancers. Nat. Genet. 53, 1334–1347 (2021).

13. Pal, B. et al. A single-cell RNA expression atlas of normal, preneoplastic and tumorigenic states in the human breast. EMBO J. 40, e107333 (2021).

14. Harris, M. A. et al. Towards targeting the breast cancer immune microenvironment. Nat. Rev. Cancer 24, 554–577 (2024).

15. Cepek, K. L. et al. Adhesion between epithelial cells and T lymphocytes mediated by E-cadherin and the alpha E beta 7 integrin. Nature 372, 190–193 (1994).

16. Svoronos, N. et al. Tumor cell-independent estrogen signaling drives disease progression through mobilization of myeloid-derived suppressor cells. Cancer Discov. 7, 72–85 (2017).

17. Rae, J. M. & Lippman, M. E. The role of estrogen receptor signaling in suppressing the immune response to cancer. J. Clin. Invest. 131, (2021).

18. Palomeque, J. Á. et al. Estrogen receptor signaling drives immune evasion and immunotherapy resistance in HR+ breast cancer. J. Clin. Invest. 136, (2026).

19. Nanda, R. et al. Effect of pembrolizumab plus neoadjuvant chemotherapy on pathologic complete response in women with early-stage breast cancer: An analysis of the ongoing phase 2 adaptively randomized I-SPY2 trial: An analysis of the ongoing phase 2 adaptively randomized I-SPY2 trial. JAMA Oncol. 6, 676–684 (2020).

20. Pusztai, L. et al. Durvalumab with olaparib and paclitaxel for high-risk HER2-negative stage II/III breast cancer: Results from the adaptively randomized I-SPY2 trial. Cancer Cell 39, 989–998.e5 (2021).

21. Cardoso, F. et al. Pembrolizumab and chemotherapy in high-risk, early-stage, ER+/HER2-breast cancer: a randomized phase 3 trial. Nat. Med. 31, 442–448 (2025).

22. De Caluwe, A. et al. First-in-human study of SBRT and adenosine pathway blockade to potentiate the benefit of immunochemotherapy in early-stage luminal B breast cancer: results of the safety run-in phase of the Neo-CheckRay trial. J. Immunother. Cancer 11, e007279 (2023).

23. Fenton, M. et al. The untapped potential of radiation and immunotherapy for hormone receptor-positive breast cancer. NPJ Breast Cancer 11, 77 (2025).

24. Darragh, L. B. & Karam, S. D. Radiation as an immune modulator: mechanisms and implications for combination with immunotherapy. Nat. Rev. Cancer 26, 270–284 (2026).

25. Ciampricotti, M., Hau, C.-S., Doornebal, C. W., Jonkers, J. & de Visser, K. E. Chemotherapy response of spontaneous mammary tumors is independent of the adaptive immune system. Nat. Med. 18, 344–6; author reply 346 (2012).

26. Guy, C. T., Cardiff, R. D. & Muller, W. J. Induction of mammary tumors by expression of polyomavirus middle T oncogene: a transgenic mouse model for metastatic disease. Mol. Cell. Biol. 12, 954–961 (1992).

27. Attalla, S., Taifour, T., Bui, T. & Muller, W. Insights from transgenic mouse models of PyMT-induced breast cancer: recapitulating human breast cancer progression in vivo. Oncogene 40, 475–491 (2021).

28. Lin, E. Y. et al. Progression to malignancy in the polyoma middle T oncoprotein mouse breast cancer model provides a reliable model for human diseases. Am. J. Pathol. 163, 2113–2126 (2003).

29. Lapidus, R. G., Nass, S. J. & Davidson, N. E. The Loss of Estrogen and Progesterone Receptor Gene Expression in Human Breast Cancer. J. Mammary Gland Biol. Neoplasia 3, 85–94 (1998).

30. Lopez-Tarruella, S. & Schiff, R. The dynamics of estrogen receptor status in breast cancer: re-shaping the paradigm. Clin. Cancer Res. 13, 6921–6925 (2007).

31. Hanker, A. B., Sudhan, D. R. & Arteaga, C. L. Overcoming endocrine resistance in breast cancer. Cancer Cell 37, 496–513 (2020).

32. Trachet, E. E., Germain, D. G., Urs, S., Draper, D. & Franklin, M. R. Abstract 920: Characterization of the orthotopic MMTV PyMT murine mammary carcinoma model following radiation and immune checkpoint blockade. in Immunology vol. 80 920–920 (American Association for Cancer Research, 2020).

33. Woodworth, M. B., Girskis, K. M. & Walsh, C. A. Building a lineage from single cells: genetic techniques for cell lineage tracking. Nat. Rev. Genet. 18, 230–244 (2017).

34. McKenna, A. & Gagnon, J. A. Recording development with single cell dynamic lineage tracing. Development 146, (2019).

35. Chen, C., Liao, Y. & Peng, G. Connecting past and present: single-cell lineage tracing. Protein Cell 13, 790–807 (2022).

36. Jones, M. G., Yang, D. & Weissman, J. S. New Tools for Lineage Tracing in Cancer In Vivo. Annu. Rev. Cancer Biol. 2023 7, 111–140 (2023).

37. Lüönd, F. et al. Distinct contributions of partial and full EMT to breast cancer malignancy. Dev. Cell 56, 3203–3221.e11 (2021).

38. Merle, C. et al. Genetic barcoding of individual cells links cancer evolutionary trajectories and prognostic outcomes. bioRxiv 2025.12.01.691488 (2025) doi:10.64898/2025.12.01.691488.

39. Kalhor, R. et al. Developmental barcoding of whole mouse via homing CRISPR. Science 361, eaat9804 (2018).

40. Leeper, K. et al. Lineage barcoding in mice with homing CRISPR. Nat. Protoc. 16, 2088–2108 (2021).

41. Spina, E. & Cowin, P. Embryonic mammary gland development. Semin. Cell Dev. Biol. 114, 83–92 (2021).

42. Kalhor, R., Mali, P. & Church, G. M. Rapidly evolving homing CRISPR barcodes. Nat. Methods 14, 195–200 (2017).

43. Serrano, A. et al. Genetic barcoding uncovers the clonal makeup of solid and liquid biopsies and their ability to capture intra-tumoral heterogeneity. Cancer Biology (2025).

44. Camargo, S. et al. Neutrophils physically interact with tumor cells to form a signaling niche promoting breast cancer aggressiveness. *Nat*. Cancer 6, 540–558 (2025).

45. Islam, M. et al. Temporal recording of mammalian development and precancer. Nature 634, 1187–1195 (2024).

46. Pfefferle, A. D. et al. Transcriptomic classification of genetically engineered mouse models of breast cancer identifies human subtype counterparts. Genome Biol. 14, R125 (2013).

47. Jiang, G. et al. Single-cell transcriptomics reveal the heterogeneity and dynamic of cancer stem-like cells during breast tumor progression. Cell Death Dis. 12, 979 (2021).

48. Yeo, S. K. et al. Single-cell RNA-sequencing reveals distinct patterns of cell state heterogeneity in mouse models of breast cancer. Elife 9, (2020).

49. Taube, J. H. et al. Core epithelial-to-mesenchymal transition interactome gene-expression signature is associated with claudin-low and metaplastic breast cancer subtypes. Proc. Natl. Acad. Sci. U. S. A. 107, 15449–15454 (2010).

50. Bartoschek, M. et al. Spatially and functionally distinct subclasses of breast cancer-associated fibroblasts revealed by single cell RNA sequencing. Nat. Commun. 9, 5150 (2018).

51. Shen, W.-K. et al. An automatic annotation tool and reference database for T cell subtypes and states at single-cell resolution. Sci. Bull. (Beijing) 70, 1659–1672 (2025).

52. Song, L. et al. TRUST4: immune repertoire reconstruction from bulk and single-cell RNA-seq data. Nat. Methods 18, 627–630 (2021).

53. Yang, Q., Safina, K. R., Nguyen, K. D. Q., Tuong, Z. K. & Borcherding, N. scRepertoire 2: Enhanced and efficient toolkit for single-cell immune profiling. PLoS Comput. Biol. 21, e1012760 (2025).

54. Gough, M. J. et al. Immune consequences of CT-guided radiation therapy of mouse mammary tumors. Journal for Immunotherapy of Cancer 1, P116 (2013).

55. Winter, E., Emiliani, F., Cook, A., Abderrahim, A. & McKenna, A. H. BASELINE: A CRISPR base editing platform for mammalian-scale single-cell lineage tracing. bioRxivorg 2025.03.19.644238 (2025) doi:10.1101/2025.03.19.644238.

56. Heyne, G. W. et al. A simple and reliable method for early pregnancy detection in inbred mice. J. Am. Assoc. Lab. Anim. Sci. 54, 368–371 (2015).

57. Bushnell, B., Rood, J. & Singer, E. BBMerge - Accurate paired shotgun read merging via overlap. PLoS One 12, e0185056 (2017).

58. Labun, K. et al. Accurate analysis of genuine CRISPR editing events with ampliCan. Genome Res. 29, 843–847 (2019).

59. Korotkevich, G. et al. Fast gene set enrichment analysis. bioRxiv 060012 (2016) doi:10.1101/060012.

