## Supplemental figures for "Mapping breast cancer lineage in radiation and immunotherapy using the REMAP mouse"

### Supplemental Figure 1

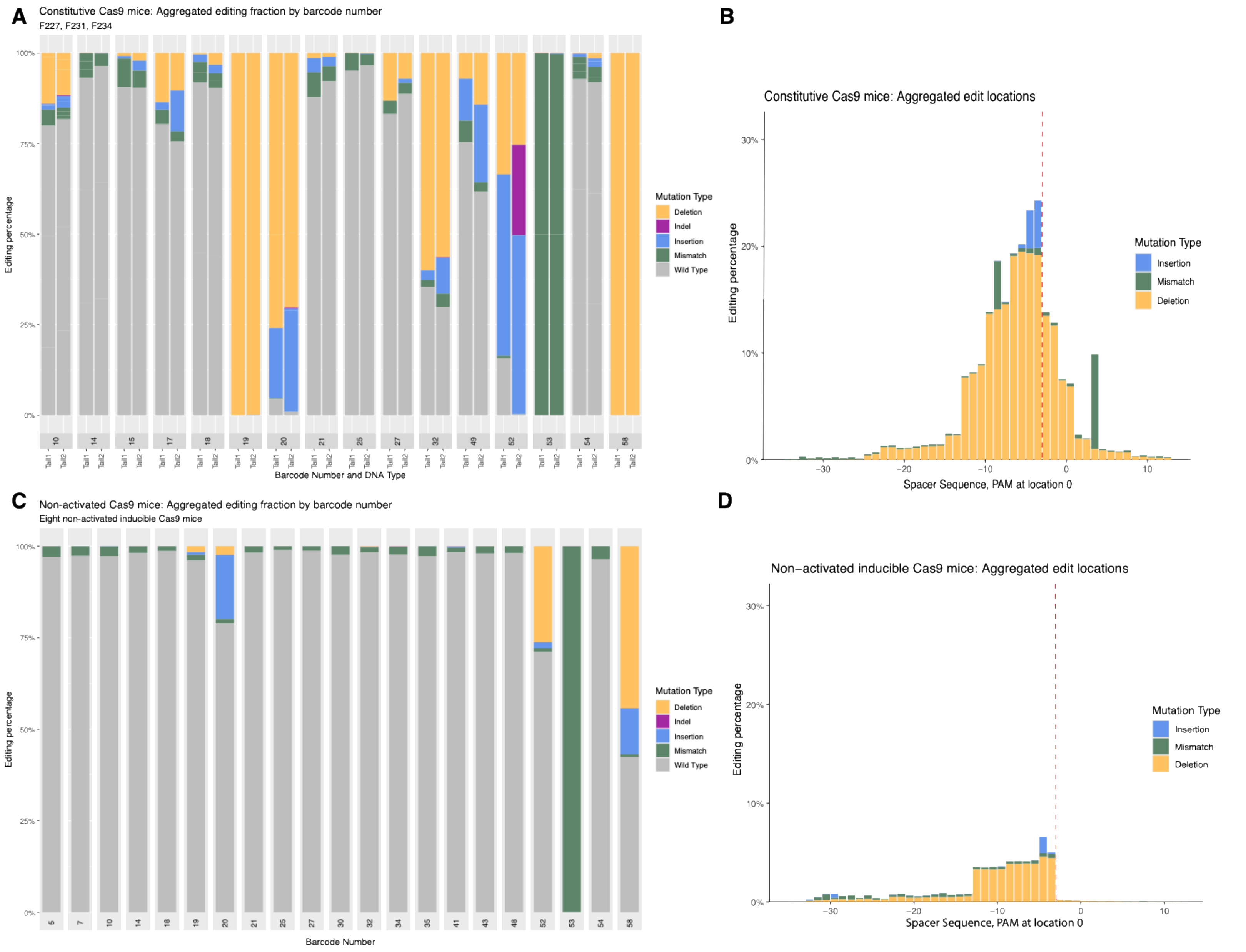

Supplemental Figure 2

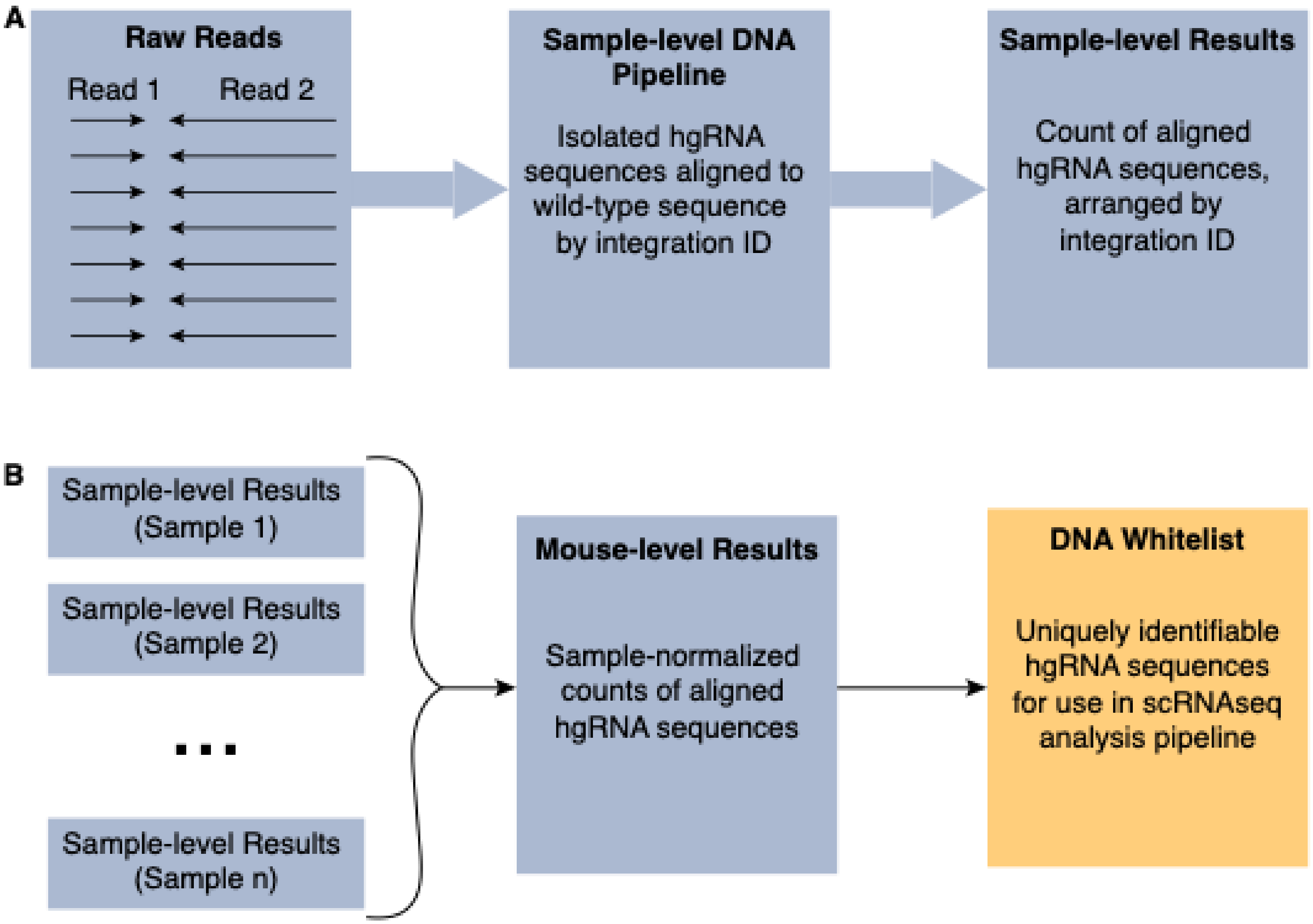

### Supplemental Figure 3

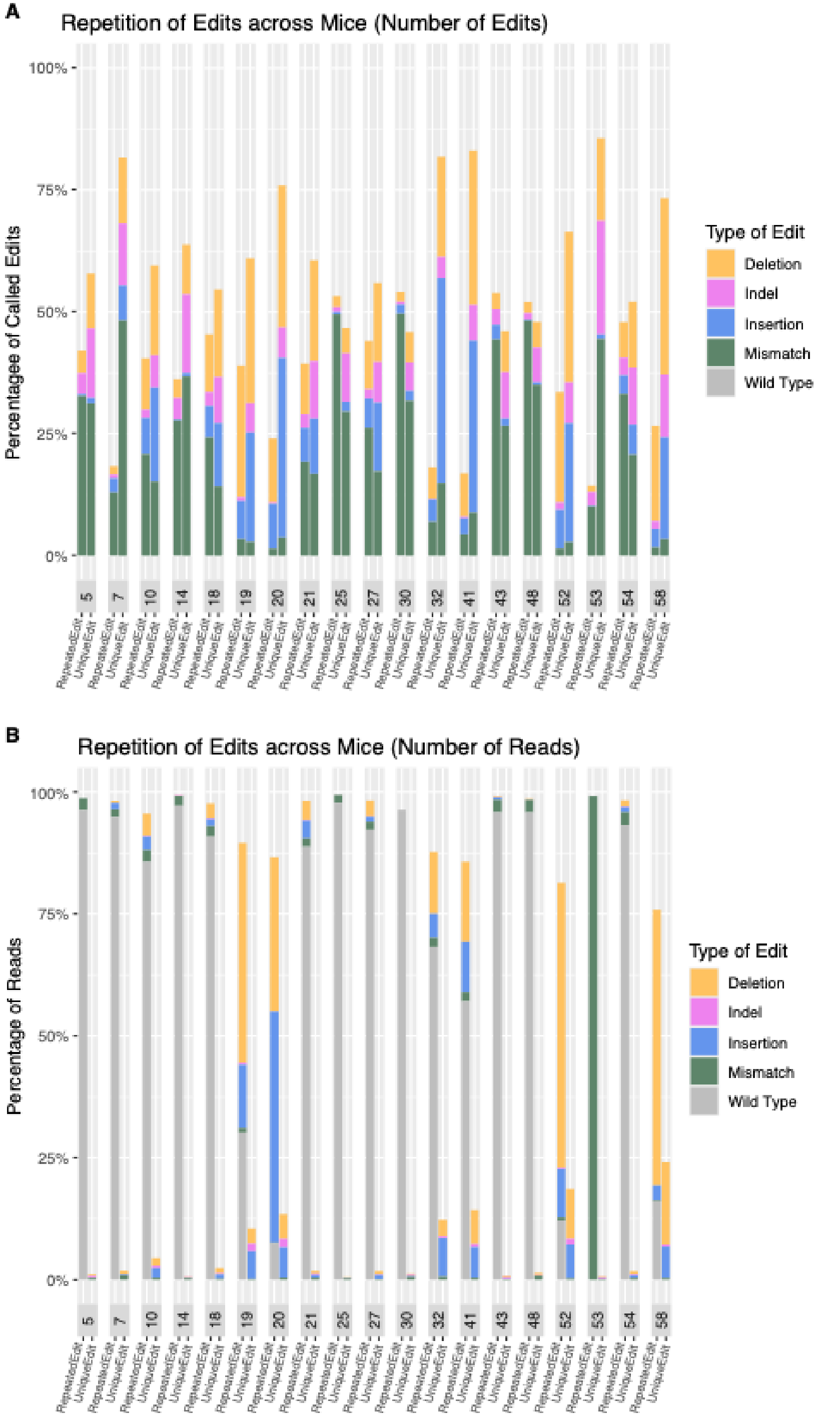

Supplemental Figure 4

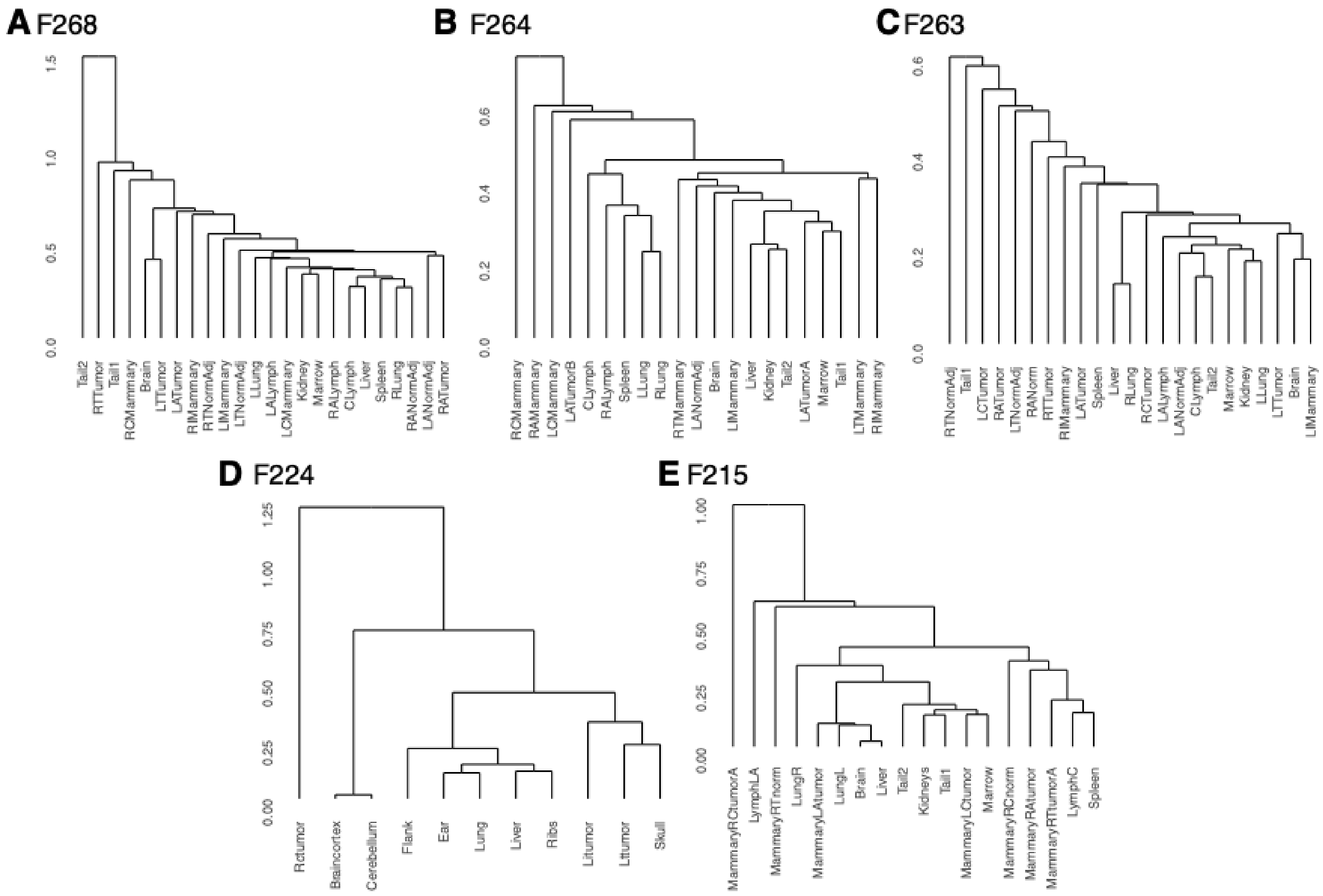

### Supplemental Figure 5

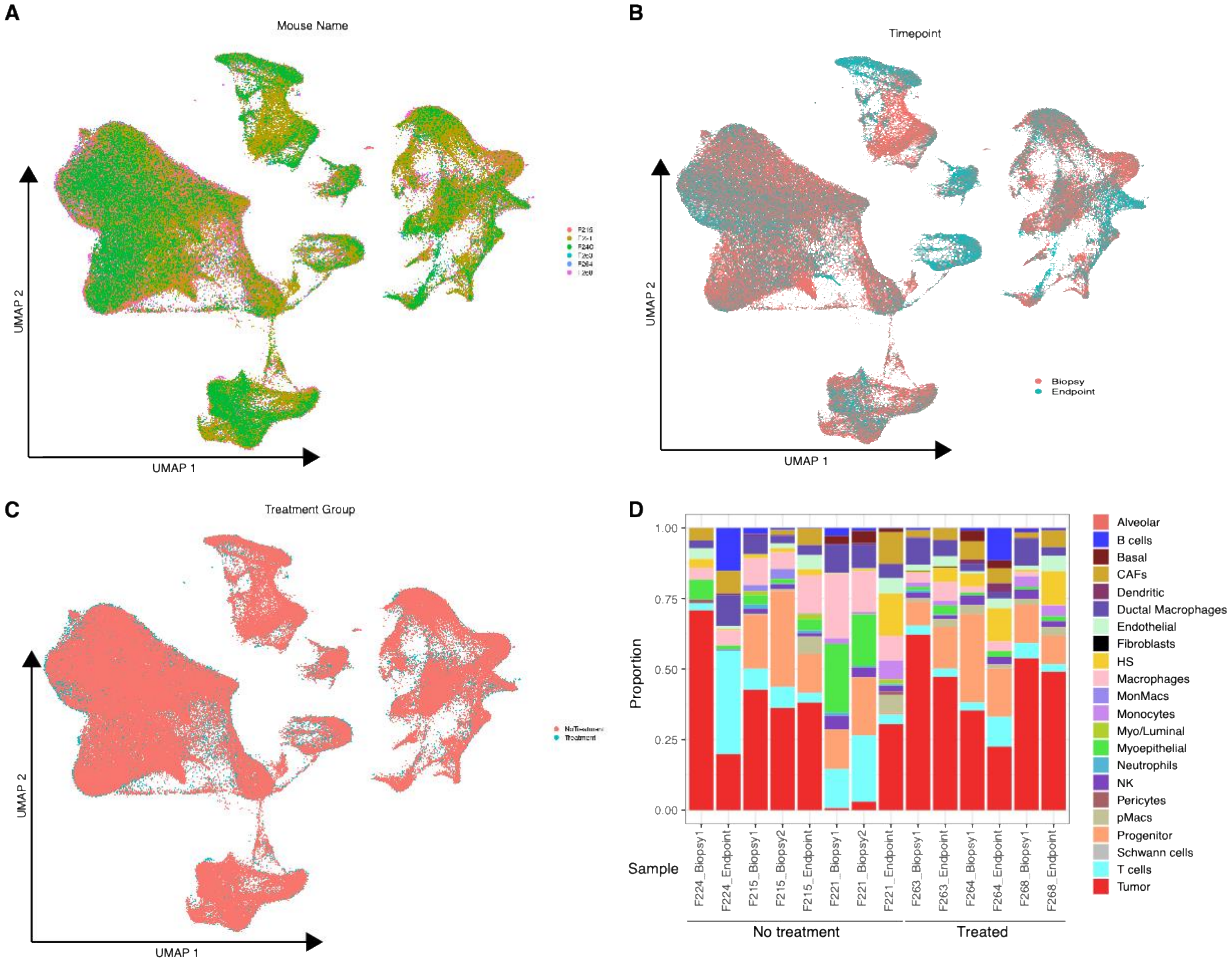

Supplemental Figure 6

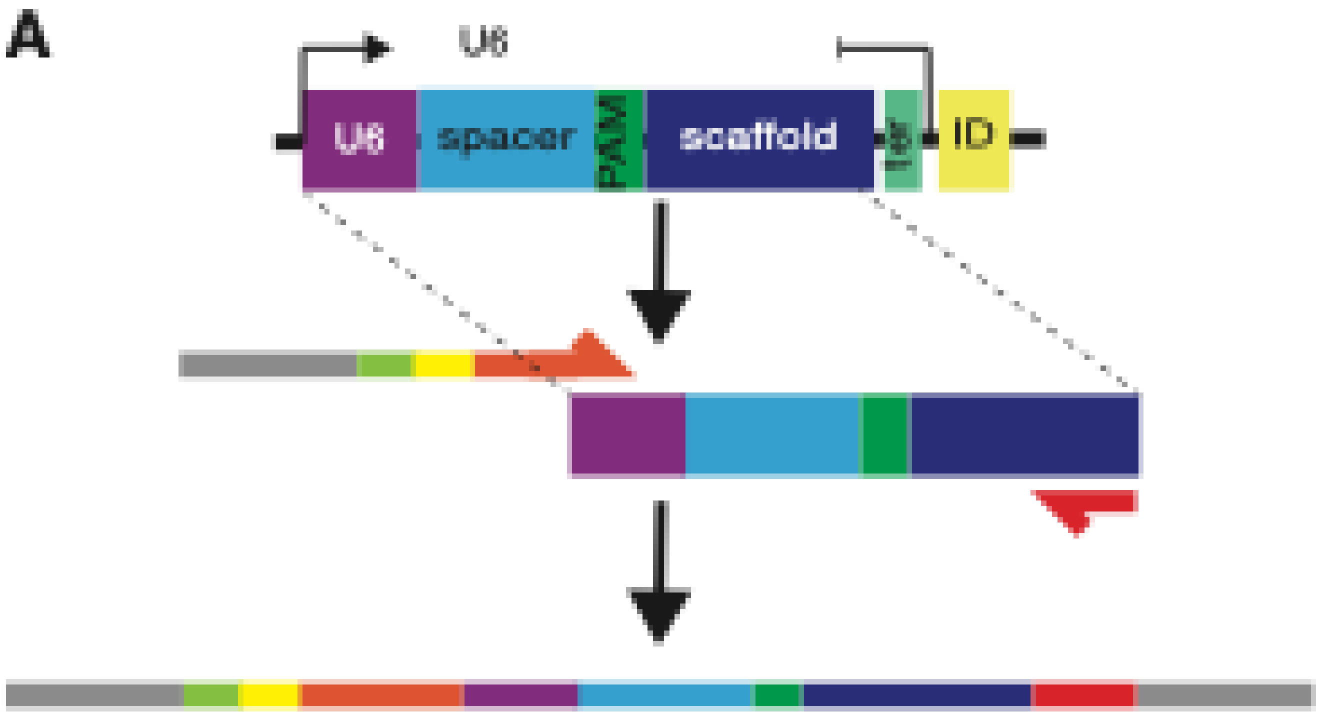

Supplemental Figure 7

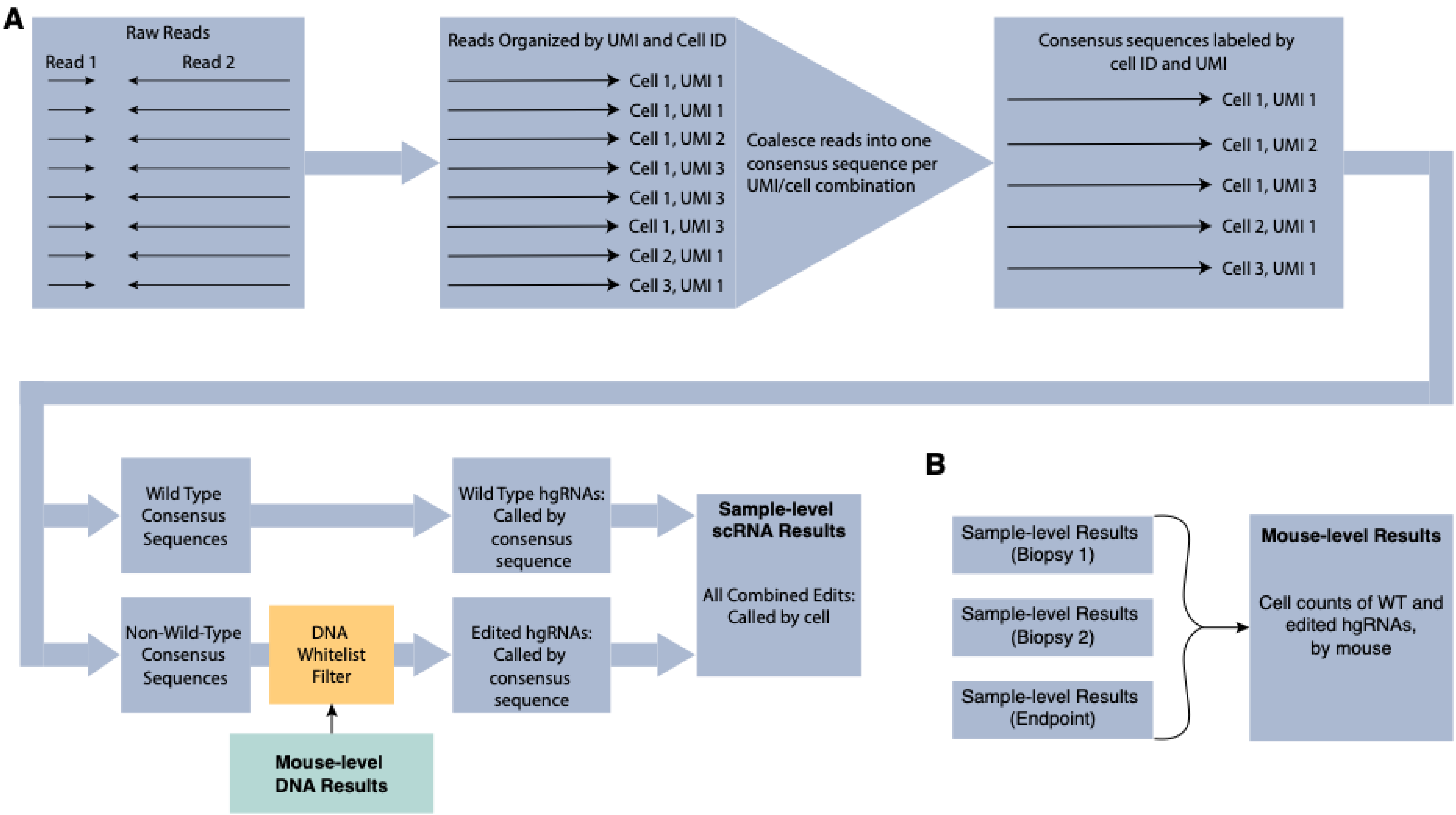

### Supplemental Figure 8

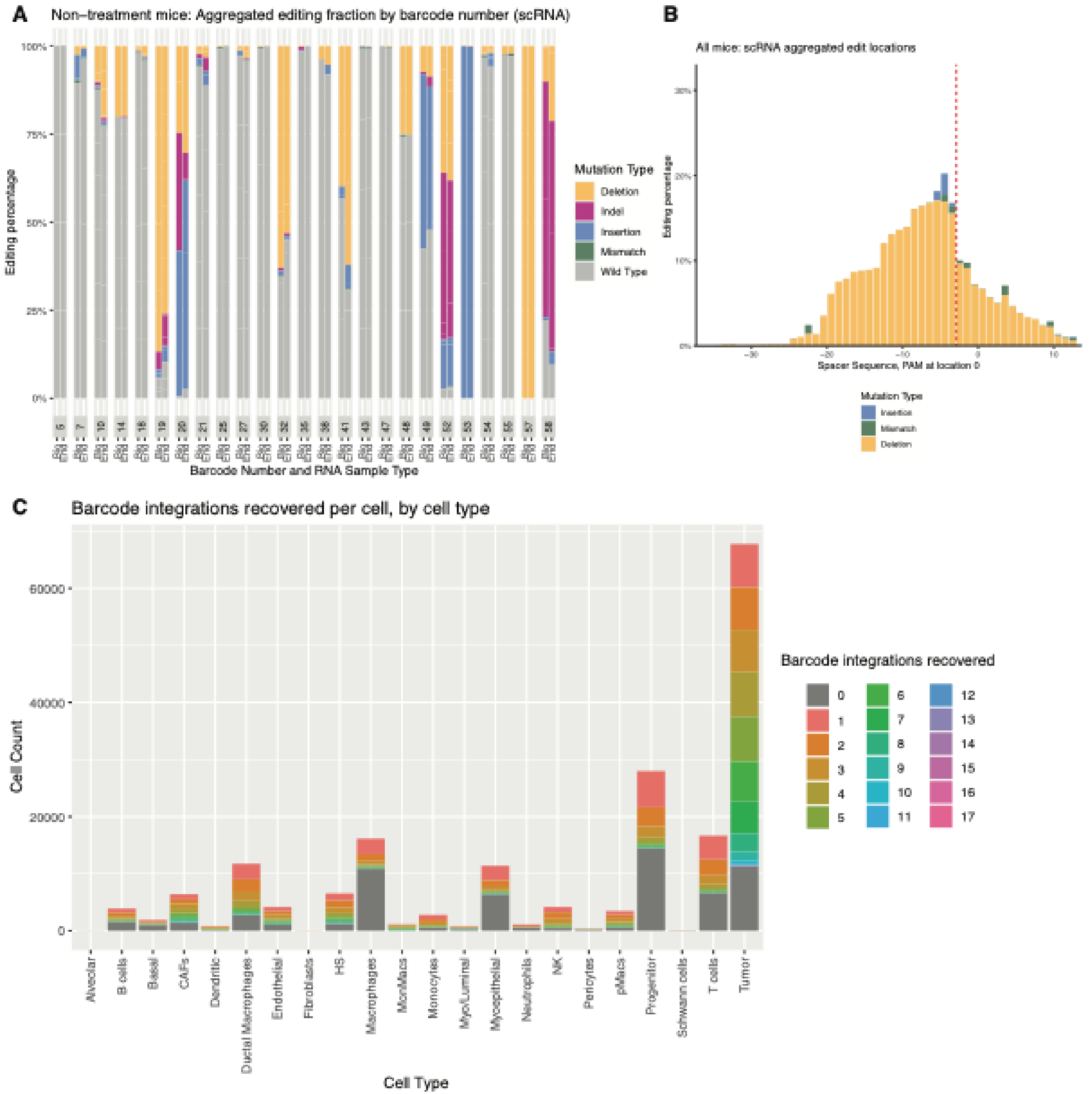

### Supplemental Figure 9

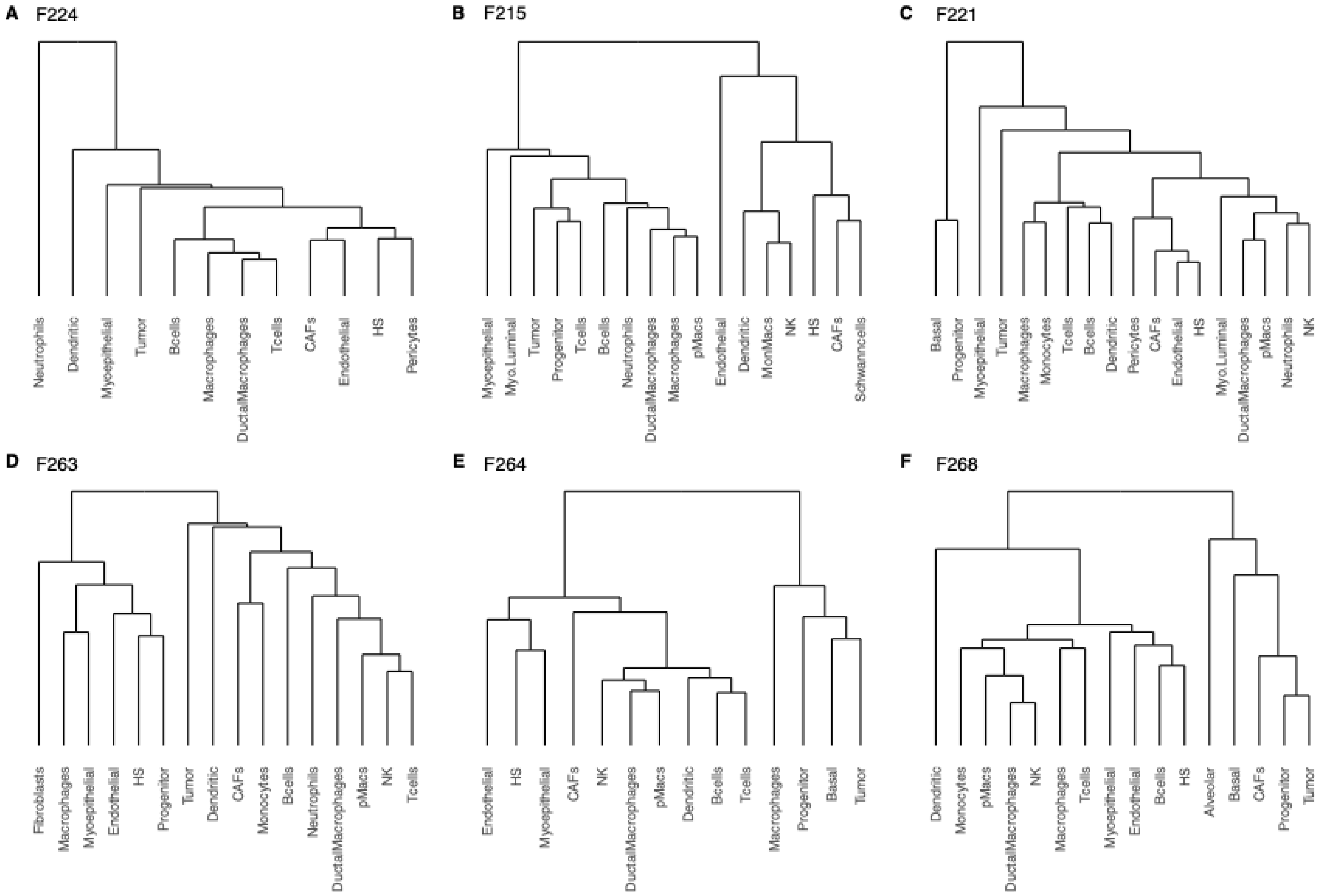

Supplemental Figure 10

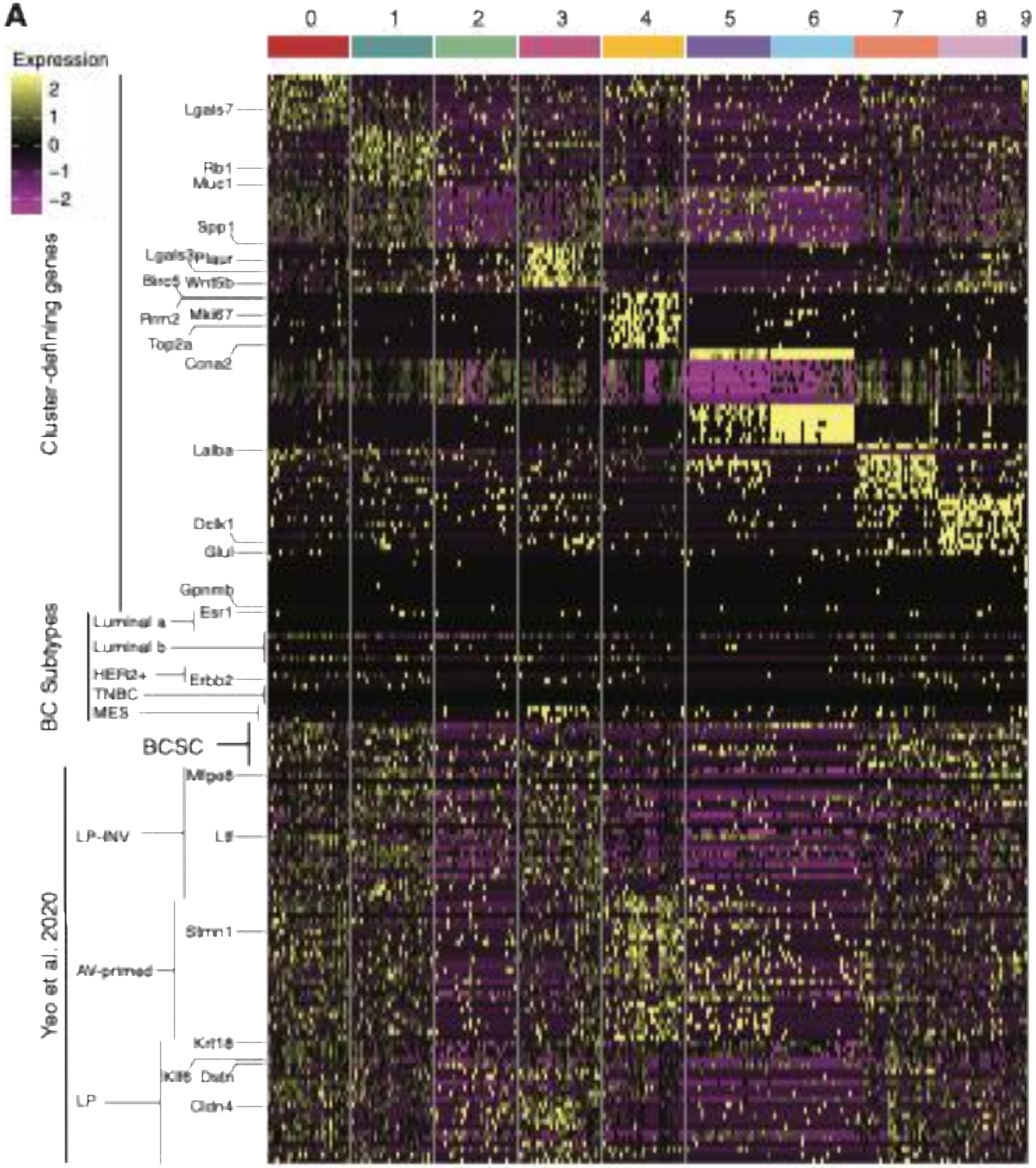

### Supplemental Figure 11

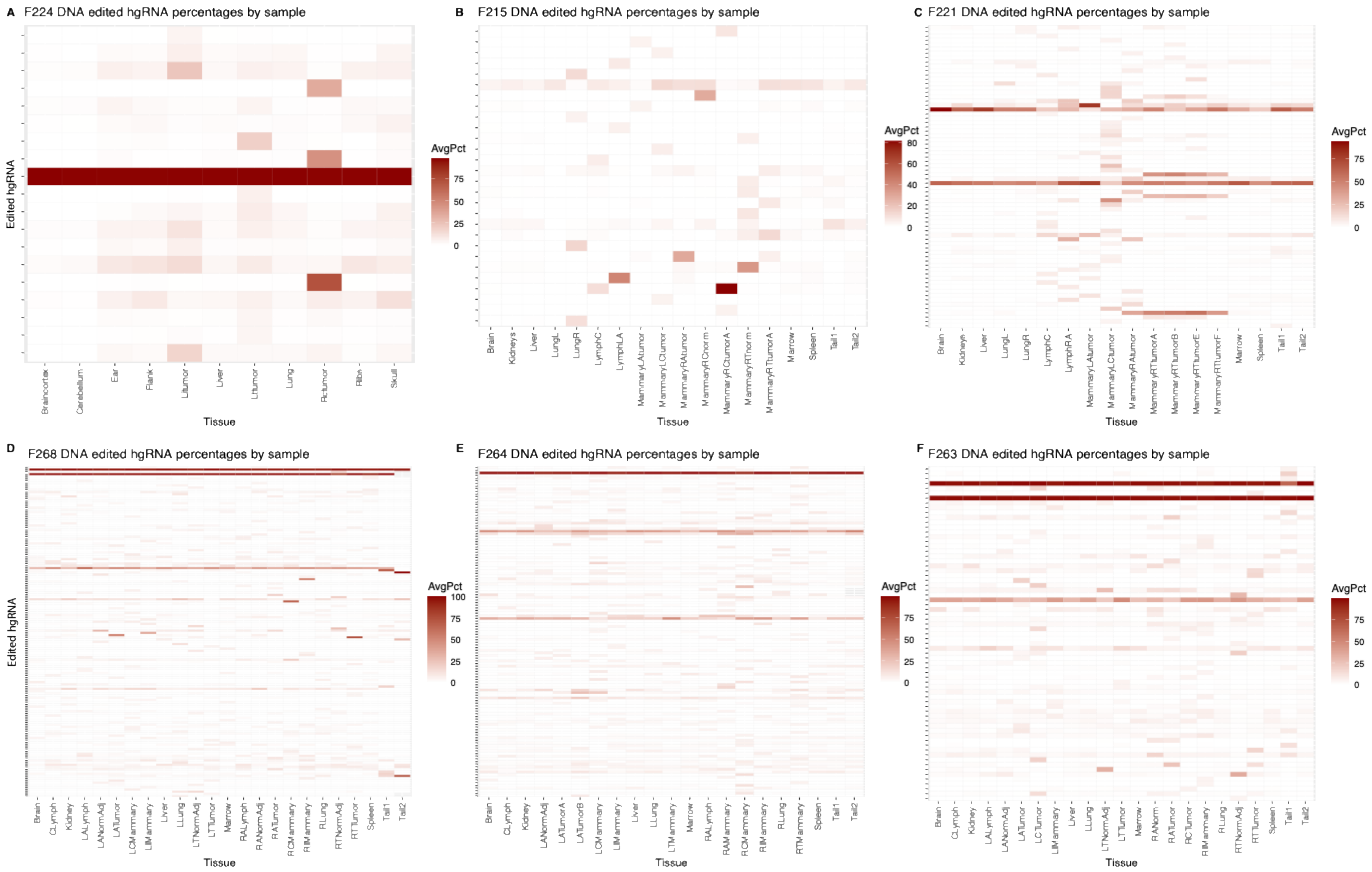

Supplemental Figure 12

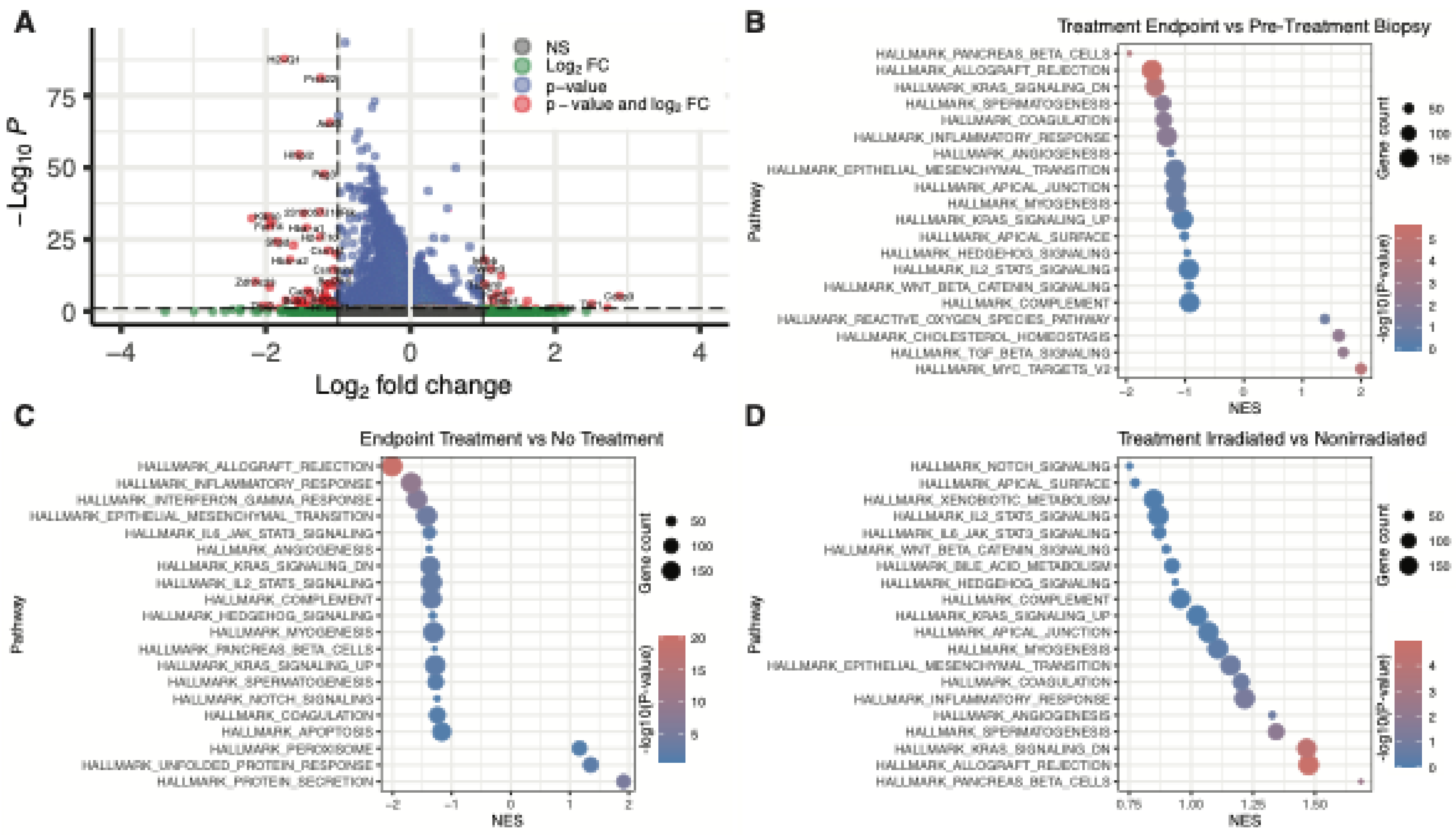

### Supplemental Figure 13

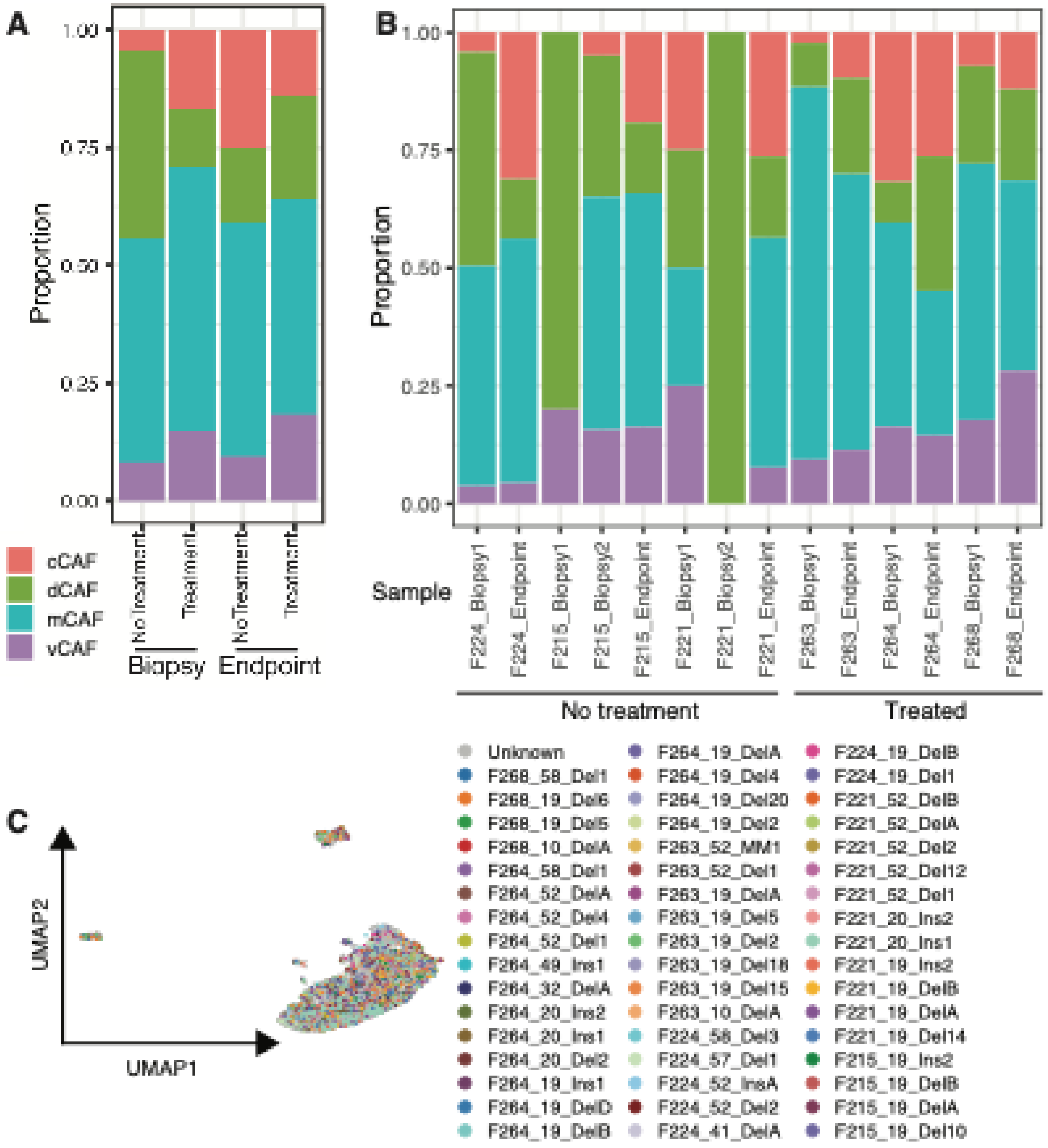

### Supplemental Figure 14

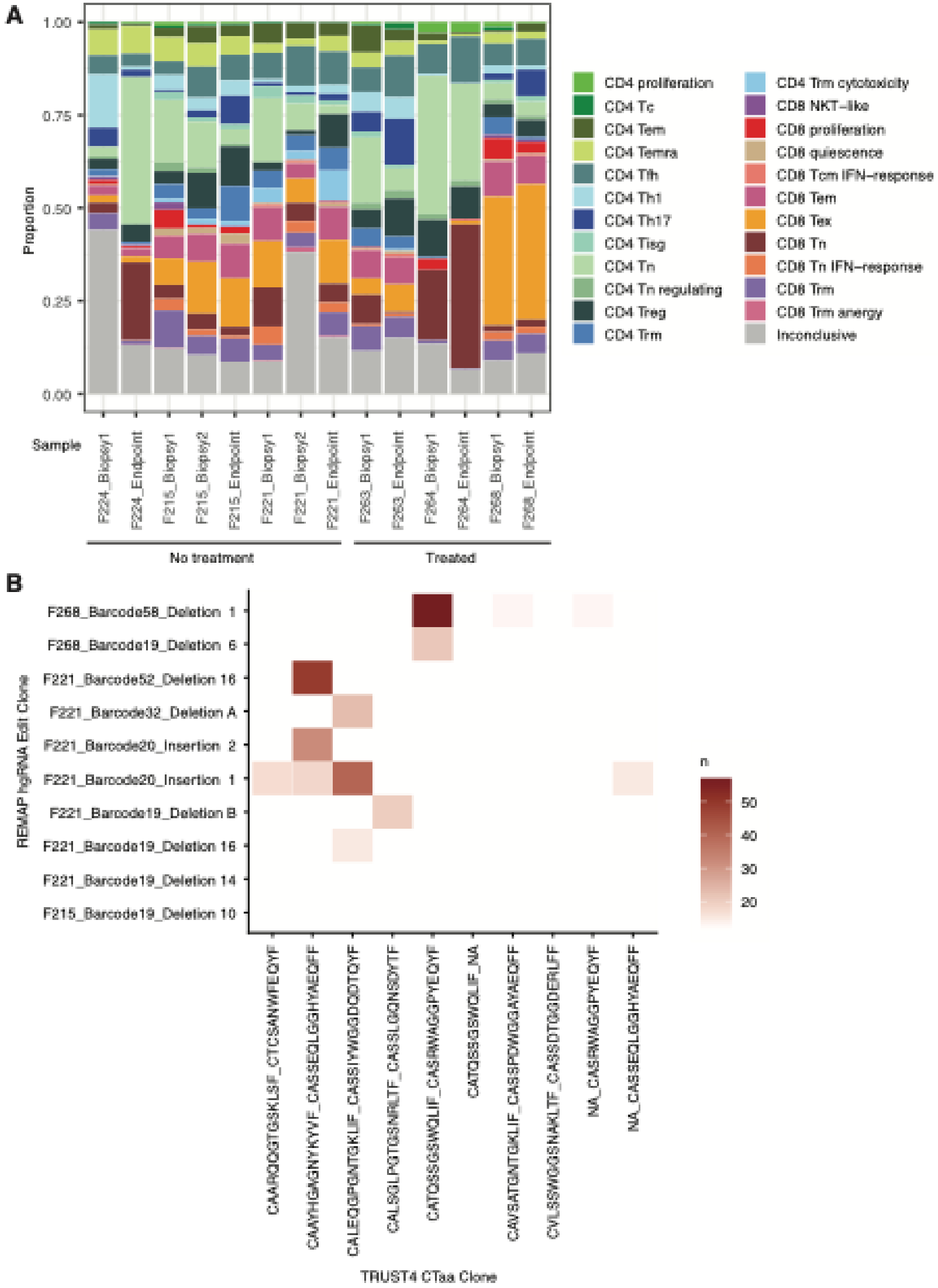
